# Microbial metabolite 4-ethylphenylsulfate (4EPS) interacts with AT1R, reduces blood pressure and outcome of AngII-induced aortic aneurysm

**DOI:** 10.1101/2025.01.29.635371

**Authors:** Terri J. Harford, Dhanachandra Khuraijam Singh, Triveni R. Pardhi, Russell Desnoyer, Tarun Ravi, Zaira Palomino Jara, Ajay Zalavadia, Kate Stenson, Sathyamangla V Naga Prasad, Sadashiva S. Karnik

## Abstract

**BACKGROUND:** Plasma accumulation of the gut microbial metabolite, 4-ethylphenylsulfate (4EPS), produced from dietary protein aromatic amino acids has been observed in correlative and associative studies of cardiovascular, renal, metabolic and neurological diseases. 4EPS level increases upon AngII infusion in mice. How 4EPS alters host physiology to contribute to progression of any disease state is currently unknown.

**METHODS:** To test the hypothesis that 4EPS interferes with angiotensin binding to AT1R, we used multiple approaches: AT1R pharmacology, cell-signaling, ex vivo vascular contraction and a mouse model of angiotensin-induced aortic aneurysm (AA) disease. ApoE-null mice were fed high-fat diet and infused with AngII, or co-infused with 4EPS and Olmesartan. BP was recorded. At the end of infusion, aortas were assessed for severity of AA, contractile response and histopathology. To evaluate signaling associated with different AA outcomes plasma proteomics analysis was done.

**RESULTS:** In vitro, 4EPS reduced the binding of angiotensin and Candesartan to AT1R and calcium signaling. Ex vivo, 4EPS decreased vasomotor response of the aorta to AngII. In vivo, 4EPS inhibited AngII-mediated increase of BP and reduced mortality from AA. Abdominal aorta remodeling in 4EPS+AngII co-infused mice showed an increase of elastin area and reduced thickening of intimal/medial layers. Plasma proteome analysis indicated significant change in actin-cytoskeletal signaling associated with reduced ERK1/2 and Filamin-A activation, and cell motility.

**CONCLUSIONS:** Benign antagonism of AT1R by 4EPS involves direct interaction with AT1R. Molecular mechanisms of 4EPS responsible for reduced AA associated mortality in mice are distinct from those of AT1R blocker, Olmesartan.

**GRAPHIC ABSTRACT:** 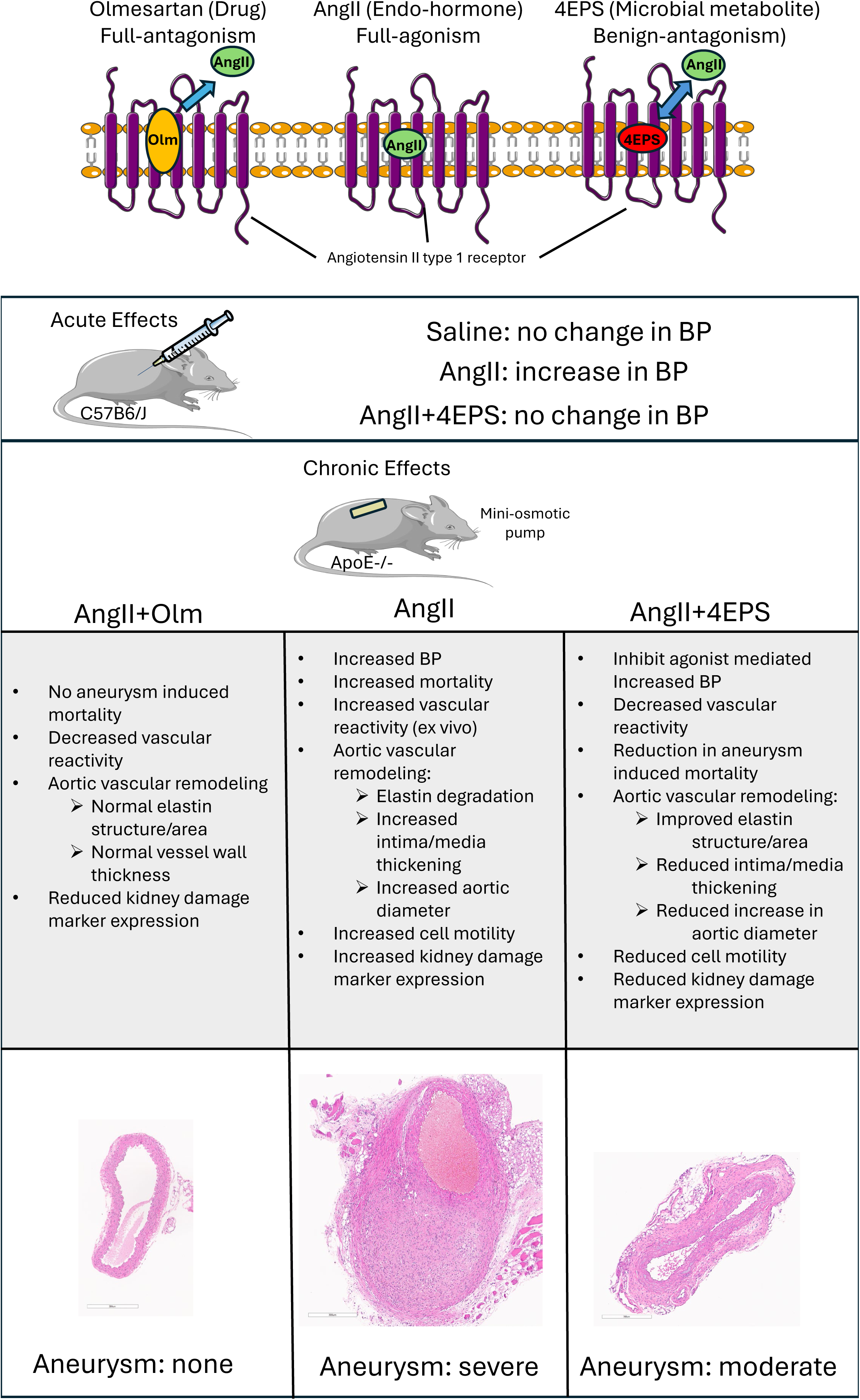

## Introduction

The renin-angiotensin system (RAS) regulates ambient changes in blood pressure (BP) and cardiovascular health through its major hormone angiotensin II (AngII). Dysregulation of AngII signaling can lead to a wide spectrum of cardiovascular diseases (CVD). Recent studies in mice reared in conventional and germ-free environment suggested that responses to AngII such as blood pressure elevation, reactive oxygen species generation, vascular inflammation, immune-cell infiltration into tissues, fibrosis of the heart and kidneys and end organ damage are subdued in germ-free mice ^1–4^. The germ-free mouse studies thus suggested that the microbiota affect the balance of benefits and risks of AngII on CVD outcomes.

Strong links between the metabolism of dietary protein, lipid and polysaccharides by gut microbial flora, and the etiology of diseases, including CVD^1,5^, obesity^6,7^, and type 2 diabetes^8^ are well established. For instance, microbial metabolism of aromatic amino acids^9–11^, branched-chain amino acids^12–14^, short-chain fatty acids^15–17^, and N-acyl amides^18^, influence blood pressure^1,19–21^, kidney functions^22^, insulin sensitivity and glucose homeostasis^23^ in mice. Gut-microbiome-induced trimethylamine-N-oxide (TMAO) production promotes development of atherosclerosis in mice^24–26^. Many human studies have examined the effects of several types of dietary components on CVDs^27^. However, our understanding of the mechanism of microbial-metabolite interactions with RAS in CVD is a knowledge gap that has greatly limited clinical translational studies.

G protein coupled receptors (GPCRs) mediate both positive and negative effects of microbial metabolites on host-health. For example, microbial metabolites, such as phenylacetylglutamine (PAGln) and phenylacetylglutamate (PAGlu), are enriched in type 2 diabetes mellitus, linked to CVD and modulate adrenergic receptor signaling in vivo^2,28–30^. Additionally, the gut microbial metabolites short-chain fatty acids^16,31^, butyrate, propionate and acetate modulate blood pressure and cardiovascular health, via GPCRs^16,32^, Gpr41^33^ and Olfr78^34,35^. A recent study showed a significant alteration in several microbial metabolites, including 4EPS in wild type C57B6 mice in response to AngII infusion^19^. 4EPS is a gut microbial metabolic product of dietary protein aromatic amino acids phenylalanine (Phe) and tyrosine (Tyr). 4EPS is significantly associated with CVD^11^, chronic kidney disease^36–38^, BP regulation^39^, and Autism Syndrome Spectrum^19,40^. Despite these important insights the molecular target and mechanisms by which 4EPS influences host physiology and function are without a mechanistic context.

As 4EPS was altered in response to infusion with AngII, key hormonal regulator of blood pressure responsible for vasoconstriction, In the present study, we tested the hypothesis that 4EPS has vaso-regulatory properties through direct interaction with angiotensin type 1 receptor (AT1R), a GPCR component of RAS. We show that 4EPS occupies the orthosteric ligand binding site of AT1R, acting as a “benign-antagonist”, reducing AngII-occupancy for AT1R and intracellular calcium mobilization, lessening ex vivo vascular contraction, lowering in vivo blood pressure and mitigating aortic aneurysm (AA) growth in a mouse model of AA. Co-administration of AngII+4EPS led to improved mean arterial pressure and reduced aneurysm formation, with no adverse effects on cardiac and renal functions. These findings have demonstrated the potential advantage of elevated circulating levels of 4EPS in ameliorating the health risks associated with hypertension. Thus, our study provides specific insights into how 4EPS regulates AngII effects on host physiology and pathology.

## Methods

### Animal Care and Use

Male and female C57BL/6J and ApoE-null (B6.129P2-*Apoe^tm1Unc^*/J Catalog 002052) were purchased from Jackson Laboratories (Bar Harbor ME). Mice were cared for in accordance with the Guide for the Care and Use of Laboratory Animals housed within the Biological Resources Unit. All mouse experimental protocols described were reviewed and approved specifically for this project by the Institutional Animal Care and Use Committee at the Cleveland Clinic. Mice were housed (4 animals per cage) with Aspen Sani-chip bedding (Teklad No. 7090A) and provided with Reverse Osmosis drinking water system. 12h light/dark cycles were maintained throughout all animal experiments with temperature controlled between 18-23°C. Mice were fed normal Harlan Teklad rodent diet with w/w (18.6% protein +44.2% carbohydrate + 6.4 fat + 0% cholesterol). In specified experiments, mice were fed high-fat (or Western) (HFD) rodent diet (TD 88137, Harlan Teklad, Madison, USA) with w/w (17.3% protein + 48.5% carbohydrate + 21.2% fat + 0.2% cholesterol and saturated fat >60%). Mice are cared for and monitored for signs of distress by a trained veterinarian and technical staff who are on duty 7 days of the week.

### Acute Ligand administration in C57BL6/J mice

12-week-old male mice (n=3), implanted with telemetry probe (model 8 HDX11) were tested for the acute responses to an IP injection of either vehicle, AngII (0.2mg/kg BW), or AngII (0.2mg/kg BW) +4EPS (0.4mg/kg BW). The systolic, diastolic and mean BP and HR were recorded for 20min pre-injection for baseline readings and 60min post-injection for induced change of readings for each mouse. Two days are given for the complete clearance of AngII and 4EPS and same treatment combination was tested three times. Body weights were recorded, and plasma, liver, and kidneys were collected from each mouse at the time of sacrifice.

### Chronic Ligand administration in AA model

ApoE-null mice fed HFD and infused with AngII were chosen as the model for experimental AA. The outcome on AA formation was monitored as described earlier^41^. This established model of AA recapitulates some key features of the human disease such as vascular inflammation, macrophage infiltration, medial elastolysis, luminal expansion and thrombus formation. Disease progression in the AngII infused group was challenged by co-infusion with either the metabolite 4EPS or the ARB, olmesartan (OLM). Male and- female ApoE-null mice at 8-weeks age were shifted to HFD and maintained for 42 days on this diet. At 10-weeks of age 4 groups (8 male and 8 female mice/group) received subcutaneous implantation of Alzet osmotic pumps loaded as follows:

1) Vehicle only (saline control group).
2) 1.4 μmol /g AngII (the endogenous hormone group).
3) 1.4 μmol /g AngII+ 2.5 μmol /g 4EPS (benign-antagonist metabolite group).
4) 1.4 μmol /g AngII+ 1.4 μmol /g OLM (ARB full-antagonist group).

Body weight for mice were recorded after recovery from surgery. At the end of 28 days of infusion body weight was recorded, mice were euthanized, and aorta and other organs collected for analysis. Osmotic pumps explanted from individual mice were examined for success of ligand infusion in each mouse.

### Blood Pressure Measurements

Blood pressure (BP) was measured in ApoE-null mice by noninvasive tail-cuff blood volume pressure recording and analysis method using the CODA multi-Channel High Throughput system, as reported earlier from our group. Mice in experimental groups were trained for 5 days to obtain consistent BP reading in the conscious state, as described earlier. BP was recorded in a blinded fashion on seventh day and day prior to osmotic pump implant surgery and on 14 and 27–28 days after for each mouse. An average of five readings were recorded for each mouse before and after, in each ligand infusion group.

### Aortic Diameter Measurement Ultrasonography

While under general isoflurane anesthesia (1-2% inhalation), mice were scanned using MS550D probe utilizing pulse wave doppler sonography allowing identification of abdominal aorta and to distinguish from vena cava. M-mode imaging was used to acquire both short and long axis images of suprarenal aortas for temporal resolution. Measurements were assessed at the maximum luminal diameter using Vevo2100 echocardiography system (Visualsonics).

### Vasomotor Function Analysis

Mouse abdominal aortas were removed, placed in cold physiological salt solution (PSS, 130mM NaCl, 4.7mM KCl, 1.18 mM KH_2_PO_4_, 1.17mM MgSO_4_-7H_2_O, 14.9mM NaHCO_3_, 5.5mM dextrose, 0.026mM CaNa (versenate), 1.6mM CaCl_2_), cleaned of perivascular tissue and cut into 2mm segments, which were mounted using 40µm platinum wire in chambers connected to DMT myography system (multi-wire myography system 620M, DMT USA, MI). Vessels were kept in PSS heated at 37°C, gassed with 5% CO_2_, 95% O_2_ and allowed to stabilize for 1 hour prior to setting resting tension. Vessels were equilibrated at a resting lumen diameter of 0.9×L100 (L100 represents vessel diameter under passive transmural pressure of 100 mm Hg). Aortic tension was displayed and recorded with LabChart software (ADINSTRUMENTS, Colorado Springs, CO). Resting tension was determined by applying increased stepwise tension from 2mN to 10mN. Fifteen minutes later, 2 potassium-induced constrictions were measured using high concentration potassium solution (74.7 mN NaCl, 60 mM KCl, 1.18 mM KH_2_PO_4_, 1.17 mM MgSO_4_-7H_2_O, 14.9 mN NaHCO_3_, 5.5 mM dextrose, 0.026 mM CaNa_2_ versenate, 1.6 mM CaCl_2_). Vessels were then pre-constricted with 1 μm phenylephrine followed by relaxation with 1 μm acetyl-β-methylcholine chloride in pre-constricted vessels to ensure intact endothelial cell layer. Arteries were used only for investigation if they constricted in response to phenylephrine and dilated in response to acetyl-β-methylcholine chloride.

### Histopathology analysis

Three to four millimeters of the supra-renal region of the aorta was dissected and fixed overnight in 1X HistoChoice tissue fixative (catalog no. H120-1L, VWR Life Science, IL). After fixation, tissues were embedded in paraffin, and transverse sections (5-μm thick) were prepared for Hematoxylin & Eosin or Verhoeff-vanGeison staining to assess how different treatments affect cell infiltration profile and elastin content. Hematoxylin & Eosin staining was automated using a Leica Multistainer ST5020 and Eosin-Phloxine staining solution (catalog no. 1082, Newcomer Supply, Middleton, WI). Samples were stained in freshly prepared Verhoeff’s Hematoxylin for 30 minutes, then differentiated for no longer than 1 minute. The remainder of the protocol was performed by standard methods. The entirety of slide samples was stained simultaneously to eliminate batch variability. Verhoeff’s Hematoxylin Solution was prepared as established by Cleveland Clinic Department of Histology and Department of Biomedical Engineering for Movat staining protocol. Slides of stained aorta section were imaged on an Aperio AT2 slide scanner (Leica Biosystems, GmbH, Wetzlar, Germany) at 20x magnification. Qualitative assessment of intima/media was performed using Hematoxylin & Eosin staining and performed using Aperio ImageScope (Leica Biosystems, GmbH, Wetzlar, Germany). Elastin in the media layer of the aorta cross section was quantified using QuPath^43^. Two separate pixel classifiers were trained in QuPath from patches of images collected to represent variations among the aorta cross sections. First one was used to detect media layer from the cross section of the aorta and a second classifier used to identify the elastin structures within the media layer. After applying the first classifier to annotate the media layer outlines were inspected and corrected as necessary. A second classifier was used to detect and measure the area of elastin structures in the media layer. Percentage of elastin in the media layer was calculated. T test was performed using GraphPad Prism version 10.2.3 for Windows, GraphPad Software, Boston, Massachusetts USA.

### Analysis of 4-ethylphenylsulfate (4EPS) in Plasma Using HPLC On-line Tandem Mass Spectrometry (LC-MS/MS)

20µl of plasma from treated mice was mixed with 80ul HPLC grade methanol and then centrifuge at 18000 rcf for 10 min. A volume of supernatant at 50µl was added into an HPLC vial and then injected onto the LC-MS/MS instrument for quantitation of 4EPS. In brief, the LC-MS/MS system used included a Vanquish UHPLC binary pump with an on-line degasser, autosampler, and column heater and a TSQ Quantiva triple quadrupole mass spectrometer with a heated electrospray (H-ESI) ion source (Thermo Fisher Scientific, Waltham, MA). The column used was a Gemini C18, 2 x 150 mm, 5μm analytical column (Phenomenex, Rancho Palos Verdes, CA). Mobile phases were A (water containing 10 mM ammonium acetate) and B (methanol containing 10 mM ammonium acetate). A gradient with a flow rate of 0.3 ml/min was used and the run started with 0% mobile phase B from 0 to 2 min. Solvent B was then increased linearly to 100% B from 2 to 6 min and held at 100% B from 6 to 12 min. The column was finally re-equilibrated with 0% B for 7 min. The HPLC eluent was directly injected into the TSQ Quantiva and the 4EPS was ionized at ESI negative mode, using Selected Reaction Monitoring (SRM) and the SRM transition (m/z) was 201 → 121 for 4EPS. Software X-Caliber was used to process the data and obtain the peak area for 4EPS. The external standard calibration curve was used to calculate the concentration of 4EPS in the plasma and urine samples.

### Proteomic analysis of murine plasma

The plasma samples were centrifuged at 20 kg for 15minutes, 5 μL plasma was taken from each sample and transferred to a new 1.5 ml Eppendorf tube. Forty-five μL 8M urea Tris-HCl pH8 lysis buffer with freshly added protease inhibitor cocktail was added into each plasma sample. Protein concentrations of the samples were determined by BCA. Forty μg of protein from each of the samples were taken based on BCA result. The samples were reduced by dithiothreitol, alkylated by iodoacetamide and precipitated by cold acetone (-20°C) overnight. Samples were centrifuged at 12000 g for 8 minutes at 0°C, and the supernatants were removed. Protein pellets were air dried then dissolved in 40μL 100mM tri-ethyl ammonium bicarbonate (TEAB) with 0.5 μg trypsin per sample. After overnight incubation, digested samples were centrifuged and 5μg of the digest from each sample was transferred to a new tube and dried down in a SpeedVac and reconstituted in 25 μl 0.1% formic acid. Ten μl sample was mixed with 10 μl 0.5x concentration iRT standards prior to LCMS analysis. To build the spectral library, 20 μg from each of 15 samples were pooled. The pooled sample was desalted using a Waters Sep-Pak C18 cartridge. The desalted pooled sample was off-line fractionated into 16 fractions using high pH reversed phase HPLC method on a Waters Xbridge C18 chromatographic column. Around 5μg peptide from each fraction was transferred to a new tube and dried down in a SpeedVac and reconstituted in 25 μl 0.1% formic acid. Ten μl sample was mixed with 10 μl 0.5x concentration iRT standards and the samples are ready for LCMS analysis. The LC-MS system was a Bruker timsTOF pro2 mass spectrometry system interfaced with a Bruker NanoElute HPLC system. The HPLC column was a BurkerFifteen 15 cm x 75 μm id reversed phase capillary chromatography column. Four μL of the sample were injected and the peptides eluted from the column by an acetonitrile/0.1% formic acid gradient at a flow rate of 0.3 μL/min were introduced into the source of the mass spectrometer on-line. The microelectrospray ion source is operated at 1.5 kV. For spectral library generation, the digest was analyzed using a data dependent acquisition (DDA) method, the instrument acquires full scans followed by MS/MS scans of the most abundant ions from the full scans in successive instrument scans. For LC-MS analysis of the samples, a data independent acquisition (DIA) method was used. The instrument acquires full scans followed by MS/MS scans of 32 fixed mass windows along the whole LC gradient. The spectral library was generated using Pulsar that is integrated in Spectronaut software package searching the DDA LC-MS data from the 16 fractions of the pooled sample. The parameters for the library generation are listed in the table below. The DIA data were analyzed using Spectronaut V17 to search against the spectral library and the mouse UniProtKB protein sequence database for the identification and quantification of proteins and peptides. False discovery rate of protein was set at 1%. The parameters for Spectronaut DIA data searching were listed in the table below.

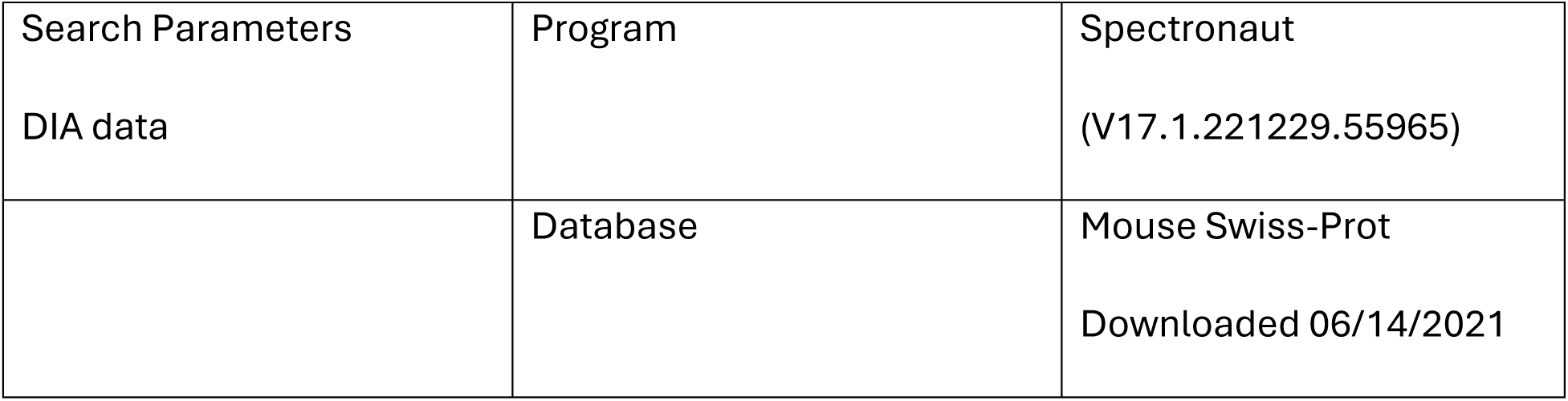
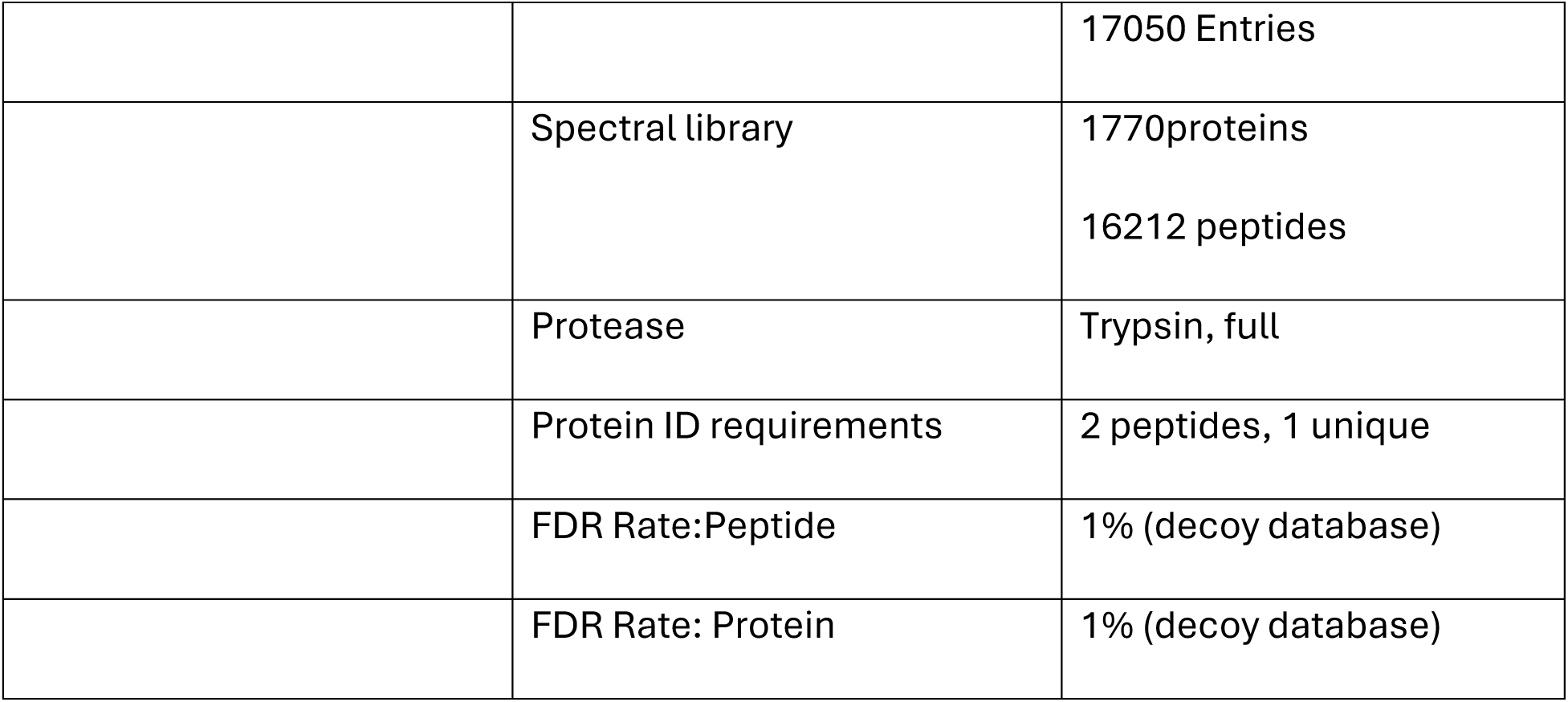

|  |  |  |
| --- | --- | --- |
| Search Parameters | Program | Spectronaut |
| DIA data |  | (V17.1.221229.55965) |
|  | Database | Mouse Swiss-Prot |
|  |  | Downloaded 06/14/2021 |
|  |  | 17050 Entries |
|  | Spectral library | 1770proteins<br>16212 peptides |
|  | Protease | Trypsin, full |
|  | Protein ID requirements | 2 peptides, 1 unique |
|  | FDR Rate:Peptide | 1% (decoy database) |
|  | FDR Rate: Protein | 1% (decoy database) |

### Necropsy

For mice that died prior to the end of each study, necropsies were performed to determine cause of death. Deaths occurring because of aortic rupture were determined by the presence of thrombus within the thoracic or retroperitoneal cavity. These mortalities were used in calculating percent survival curves.

### RT-qPCR

Total RNA was isolated from the cells using Qiagen RNeasy mini kit (Qiagen, Hilden, GE) according to the manufacturer’s instructions, and 500ng of RNA was used in a 20-µL SuperScript III RT (Invitrogen, Carlsbad, CA) reverse transcription reaction. One microliter of this reaction was used for qPCR, with 0.3 µM each of primer and SsoAdvancedSYBR Green Supermix (Bio-Rad, Hercules, CA). Transcript expression was normalized using glyceraldehyde 3-phosphate dehydrogenase (GAPDH) as the housekeeping gene. The relative change in gene expression was calculated using the following formula: % change =2^-(ΔΔCq) = 2[ΔCq (treated samples) - ΔCq (control samples)]^, where ΔCq = Cq (gene of interest) – Cq (GAPDH housekeeping gene) and Cq was the threshold number. Vehicle was used to define 100% baseline for comparison purposes. All primers used were intron-spanning and each sample was run in triplicate.

Primers:

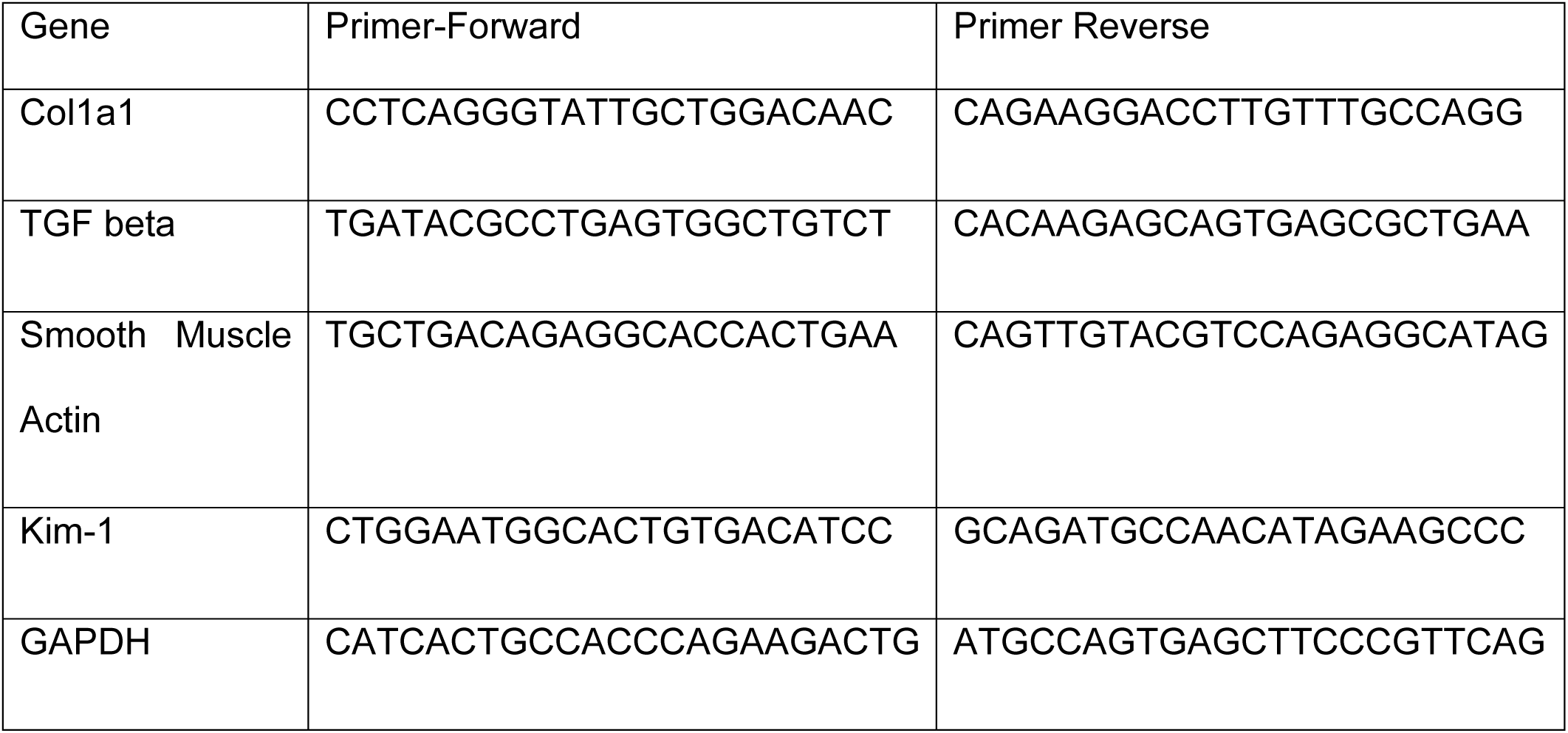

### Reagents

Angiotensin II, [Sar1]-AngII, (Bachem, Torrance CA).^125^I-Angiotensin IV (purchased from Dr. Robert Speth, Georgetown University), olmesartan medoxomil (Daichi Sankyo Chemical Pharma), 4-ethylphenyl Sulfate (4EPS, ApexBio catalog B6051), 4-(tert-butyl)phenyl hydrogen sulfate (4EPS_S4), 4-benzylphenyl hydrogen sulfate (4EPS_S40) and [1,1’-biphenyl]-4-yl hydrogen sulfate (4EPS_S46) (Chemhere, Hong Kong, China)

### Cell Culture

HEK293 (Human embryonic kidney cells) stably expressing rat AT1aR protein were grown in Dulbecco’s Modified Eagle Media (DMEM, Invitrogen) supplemented with 10% fetal bovine serum and G418 (1mg/ml). <u>Mo</u>use <u>V</u>ascular <u>A</u>ortic <u>S</u>mooth muscle cells were acquired from the ATCC (cat # ATCC® CRL-2797™). These cells are immortalized aortic smooth muscle cells from mice, using SV40 large T antigen. MOVAS cells obtained from ATCC and cultured in DMEM cells were tested for AT1R functional response using calcium assay. Cells did not show measurable Ca^2+^ response. MOVAS cells were then transfected with pcDNA-HA-AT1aR expression plasmid that contained hygromycin resistance. The MOVAS-AT1R clone stably expressing HA-AT1R as determined by FACS analysis using the anti-HA-Alexa Fluor 488 (Cell Signaling Technology, Danvers, MA catalog 349 #2350S) were grown in DMEM containing 0.2mg/mL G418 and 200μg/mL hygromycin. During experiments, serum was withdrawn for 3 hours prior to treatment/assessment as indicated.

### Cell Viability Assay

MOVAS cells plated at equal density were treated overnight in DMEM with serial dilution of 4EPS or AngII (range 1nM-100µM) or H_2_O_2_ (50mM 3 hours) prior to addition of CellTiter-Glo 2.0 reagent (Promega Corporation, Madison WI). Readings were taken using FlexStation 3 Multimodal Microplate Reader (Molecular Devices, San Jose CA).

### Calcium Assay

Calcium levels were measured using the Fluorescent Imaging Plate Reader Calcium 5 Assay kit (Molecular Devices) as described previously^42^. HEK-AT1R or MOVAS-AT1R cells were seeded in a 96-well clear bottom black plate. Following serum starvation of cells, calcium-sensitive dye was added. FlexStation 3 was programmed to add ligands at concentrations indicated to the cells and monitor the fluorescence before and after the addition of ligands. Changes in intracellular calcium were recorded by measuring ΔF/F (max-min) and are represented as relative fluorescence units. The dose–response curves were calculated assuming maximum stimulation by AngII. Data were analyzed using the sigmoidal dose–response function built into GraphPad Prism 10.

### Western Blot

Protein lysates were prepared by lysis in M-PER buffer (ThermoFisher, catalog 78501) supplemented with Halt protease/phosphatase inhibitors (Thermofisher catalog 78440). Equal amount of protein was separated by SDS-PAGE then transferred to nitrocellulose membranes. After blocking (LICORbio, Intercept blocking buffer catalog 927-700001), membranes were incubated with primary antibodies: anti-FlnA (Cell Signaling 4762), anti-phosphor FlnA (Cell Signaling 4761), anti-ERK (p42/44) (Cell Signaling 4696), anti-phosphor ERK (Cell Signaling 4370) and beta actin (Millipore). IRDye800CW conjugated goat anti rabbit and IRDye700CW conjugated donkey anti mouse secondary antibodies were applied, and signals were visualized using Odyssey DLx imaging system (LICORbio) scanning both 700and 800nm. Band intensities were quantified using ImageStudio Lite and presented as ratios of phosphorylatedto total protein.

### Radioactive Ligand Binding Assay

Ligand binding was analyzed using membranes prepared from HEK-AT1R as described earlier and suspended in membrane binding buffer (140 mM NaCl, 5.4 mM KCl, 1 mM EDTA, 0.006% bovine serum albumin, 25 mM HEPES, pH 7.4). 10μg of homogeneous cell membrane were used per well. Competition binding assays were performed under equilibrium conditions, using radioligand [^3^H]-Candesartan (4nM) and 0.04-1000nM concentrations of [Sar1] Ang II with or without 1mM 4EPS or analogs of 4EPS at a range of 1mM to 1nM. [^3^H]-Candesartan was a gift from Astra-Zeneca. Binding reaction was terminated by filtering the binding mixture through Brandel GF/C glass fiber filters, washed with buffer (20 mM sodium phosphate, 100 mM NaCl, 10 mM MgCl_2_, 1 mM EGTA, pH 7.2). The bound ligand concentration was determined as the counts/min (MicroBeta2 Plate Counter, PerkinElmer Life Sciences). Nonspecific binding was measured in the presence of [Sar1] Ang II. The filter membranes were soaked in 7 ml of Ecoscint A scintillation fluid (National Diagnostics) and incubated for 1 h at room temperature. The bound ligand fraction was determined as the disintegrations/min using a Beckman LS 6000 Liquid Scintillation Counter (Global Medical Instrumentation). Two independent experiments, each done in triplicate, were performed and the resulting values were pooled to a mean curve which is displayed. The binding kinetics were analyzed by the nonlinear curve-fitting program GraphPad Prism 8. The means ± SE for the LogIC50 values were calculated as described earlier.

### Effect on saturation binding in response to 4EPS

Saturation binding assays with ^125^I-AngIV. were performed under equilibrium conditions, with ^125^I-AngIV concentrations ranging between 1.6 and 27nM (specific activity, 16 Ci/mmol) as duplicates in 96-well plates for 1hr at room temperature in the presence of 1mM 4EPS or 4EPS analogs (S4, S40 or S46). The binding kinetics were analyzed by nonlinear curve-fitting program GraphPad Prism 8, which yielded the mean ± SE for the Kd and Bmax values.

### Protein Preparation

The initial coordinates of AT1R, bound with Olmesartan and AngII, were retrieved from the Protein Data Bank (PDB IDs: 4ZUD and 6OS0) for subsequent Glide docking calculations (Glide, Schrödinger, LLC, NY, USA). The protein was prepared by applying energy minimization through the Protein Preparation Wizard, using the OPLS4 (Optimized Potentials for Liquid Simulations) force field ^44^. Missing loop segments were completed using Prime (Schrödinger, LLC) ^45^. During protein preparation, progressively weaker restraints were applied to non-hydrogen atoms, with structure refinement performed according to the default Schrödinger protocol. Glide, which uses the full OPLS4 force field at an intermediate docking stage, is considered more sensitive to geometric details compared to other docking tools. The most likely positions of hydroxyl and thiol hydrogen atoms, protonation states, tautomers of His residues, and Chi “flip” assignments for Asn, Gln, and His residues were determined. Final energy minimization was applied to the protein until the root-mean-square deviation of non-hydrogen atoms averaged 0.3 Å.

### Ligand Preparation

All the ligands were prepared using LigPrep package from Schrodinger, LLC, NY, USA (LigPrep, 2018) by assigning appropriate bond order. All the ligands were converted to mae format (Maestro, Schrodinger, LLC, NY, USA), geometrically optimized and partial atomic charges were computed. Then, at most, 32 poses were generated with different steric features for the subsequent docking study (Schrödinger Release 2024-3: LigPrep, Schrödinger, LLC, New York, NY, 2024.)

### Glide XP Docking

Rigid docking of all ligands was performed using the Glide XP module (Schrödinger, LLC). The shape and properties of the receptors were represented on a grid through several sets of fields that progressively provided more accurate scoring of ligand poses. The experiments were conducted using default parameters. Ligands were docked into the active site using the ’extra precision’ (XP) Glide algorithm (Schrödinger Release 2024-3: LigPrep, Schrödinger, LLC, New York, NY, 2024.) ^46–48^.

### Induced Fit Docking (IFD)

Induced Fit Docking (IFD) (Schrödinger, LLC, NY, USA) was used to dock all the ligands into AngII and Olmesartan-bound pocket of AT1R. Initially, ligands were docked into a rigid receptor model with scaled-down van der Waals (vdW) radii. A vdW scaling factor of 0.5 was applied to both protein and ligand nonpolar atoms. A constrained energy minimization was performed on the protein structure, keeping it close to the original crystal structure while resolving any steric clashes. This minimization used the OPLS4 force field with an implicit solvation model. Glide XP mode was employed for the initial docking, and ligand poses were retained for subsequent protein structural refinements. Prime (Schrödinger, LLC, NY, USA) was then used to generate induced-fit protein-ligand complexes, where both sidechain and backbone refinements were carried out Schrödinger Release 2024-3: Prime, Schrödinger, LLC, New York, NY, 2024.. All residues with at least one atom located within 4.0 Å of the corresponding ligand pose were included in the Prime refinement. The refined complexes were ranked by Prime energy, and the top 20 receptor structures within 30 kcal/mol of the lowest energy structure were selected for a final round of Glide docking and scoring. In the final step, each ligand was re-docked into the top 20 refined structures using Glide XP^47^.

### Statistical analysis

Data are summarized as means ± SEM. Normality of the distribution of data was confirmed using the D’Agostino-Pearson normality test. Student t-test was used when comparing 2 variables. Comparison of more than 2 variables was performed with 1-way ANOVA with Dunnett’s multiple comparison test when normality test passed. If normality was not achieved, one way ANOVA was performed using Kruskal Wallace followed by Dunnett’s multiple comparison. Dose response curves were done using nonlinear regression and normality was also tested using D’Agostino-Pearson normality test. Radiotelemetry data analysis was based on assessing area under the curves by one way ANOVA. Tail cuff blood pressure was analyzed by one way ANOVA followed by Tukey test. All statistical analysis was performed using GraphPad Prism 10.

## Results

### Effect of 4EPS on AngII interaction with AT1R

Cheema and Pluznick previously reported that multiple microbial metabolites are altered in the plasma metabolome of C57B6/J mice infused with AngII^19^. We investigated direct influence of six significantly altered metabolites shown in **Fig 1** on activation of AT1R by AngII. HEK293 cells stably expressing rat AT1aR (HEK-AT1R, ∼5pmol/mg) were used to measure changes in AngII-dependent intracellular Ca^2+^ response. The dose of the selected metabolites was chosen based on reported clinical values in disease states or, if not reported, 10-fold increase found in human plasma in non-disease states. The effect on AT1R activation by AngII was insignificant upon pretreatment with five different metabolites. In contrast, Ca^2+^ response was significantly inhibited ∼10-fold by 4EPS [**Fig 1**] when compared to the vehicle treated cells (p<0.0001). In HEK cells not expressing the rat AT1aR, AngII and the metabolites did not elicit a Ca^2+^ response. The inhibitory effect of 4EPS was confirmed to be not due to any cytotoxic effects (see **Fig S1**). We therefore chose 4EPS for further investigation.

**Figure 1:**
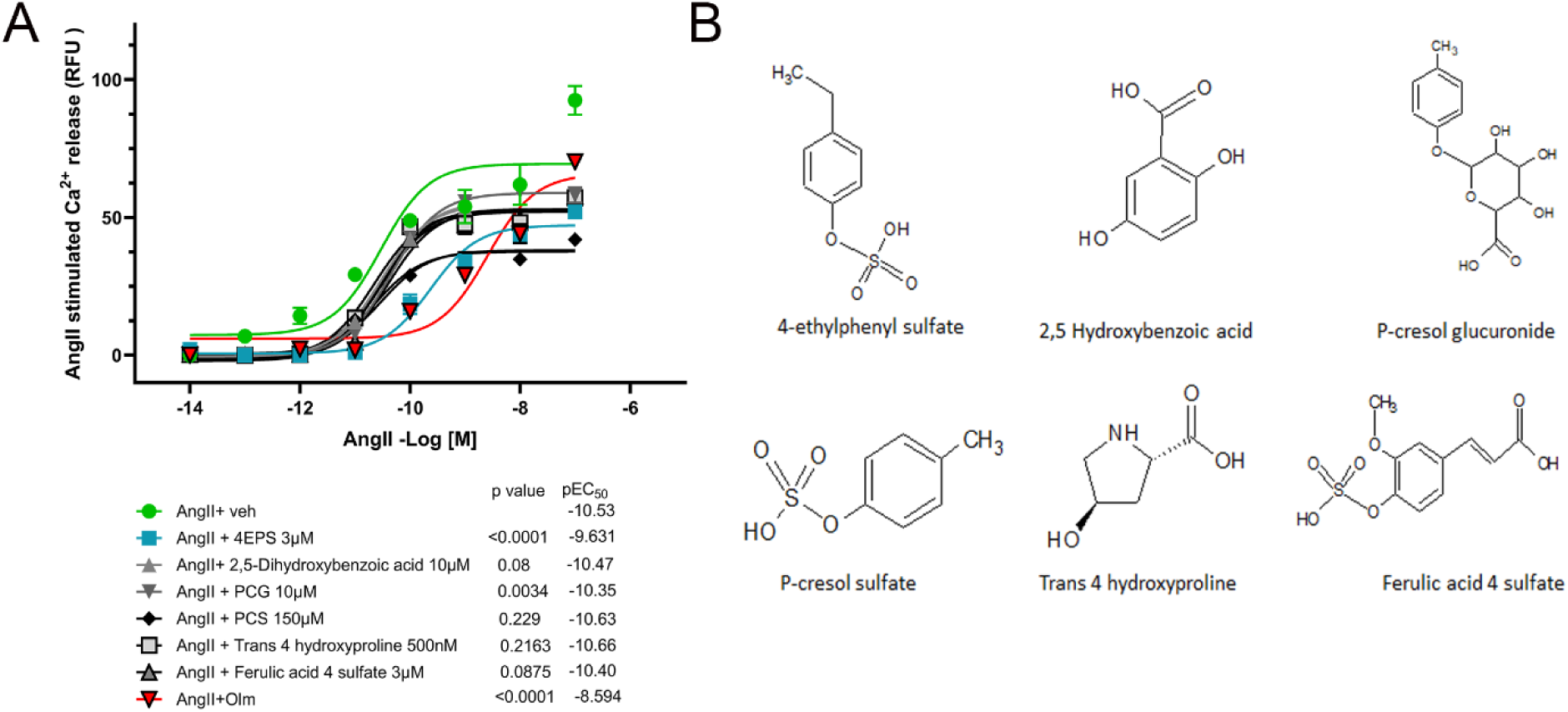
Vasoactive property of microbial metabolites. Effect of microbially derived metabolites on AngII-mediated signaling in HEK293 expressing AT1R cells. A. HEK293-AT1R cells were preincubated with vehicle (normal saline), Olmesartan (100nM) or metabolites, incubated with calcium 5 dye then stimulated with AngII dose response. Calcium mobilization was monitored using FlexStation 3. Data expressed as normalized max-min. Analysis performed using GraphPad Prism 10 non-linear regression comparison logEC50 AngII only as hypothetical value to determine p-value. (table below). N=3. B. Structures of bacterially derived metabolites assessed.

To test the hypothesis that interference with AngII binding is the basis of 4EPS antagonism of AT1R function, we performed ligand binding studies using the partial agonist ^125^I-AngIV. Total membrane isolated from mouse aortic smooth muscle cells (MOVAS) stably expressing AT1aR were used for saturation binding assays. Reduction of ^125^I-AngIV binding in the presence of 4EPS (1mM) was significant (*p<0.05, ****p<0.001 student t-test) as seen in **Fig 2A**. Competition binding assays were performed. The antagonist tracer ^3^[H]-Candesartan displacement by Sar1-AngII +/- 4EPS showed an increase of the inhibitory concentration (IC_50_) values in the presence of 4EPS [**Fig 2B**]. However, the change in IC_50_ did not reach significance (p=0.07). The reasons may be technical, i.e. [Sar1]AngII binding is stronger than AngIV and also higher receptor density in the membranes may mask the inhibitory effect of 4EPS.

**Figure 2:**
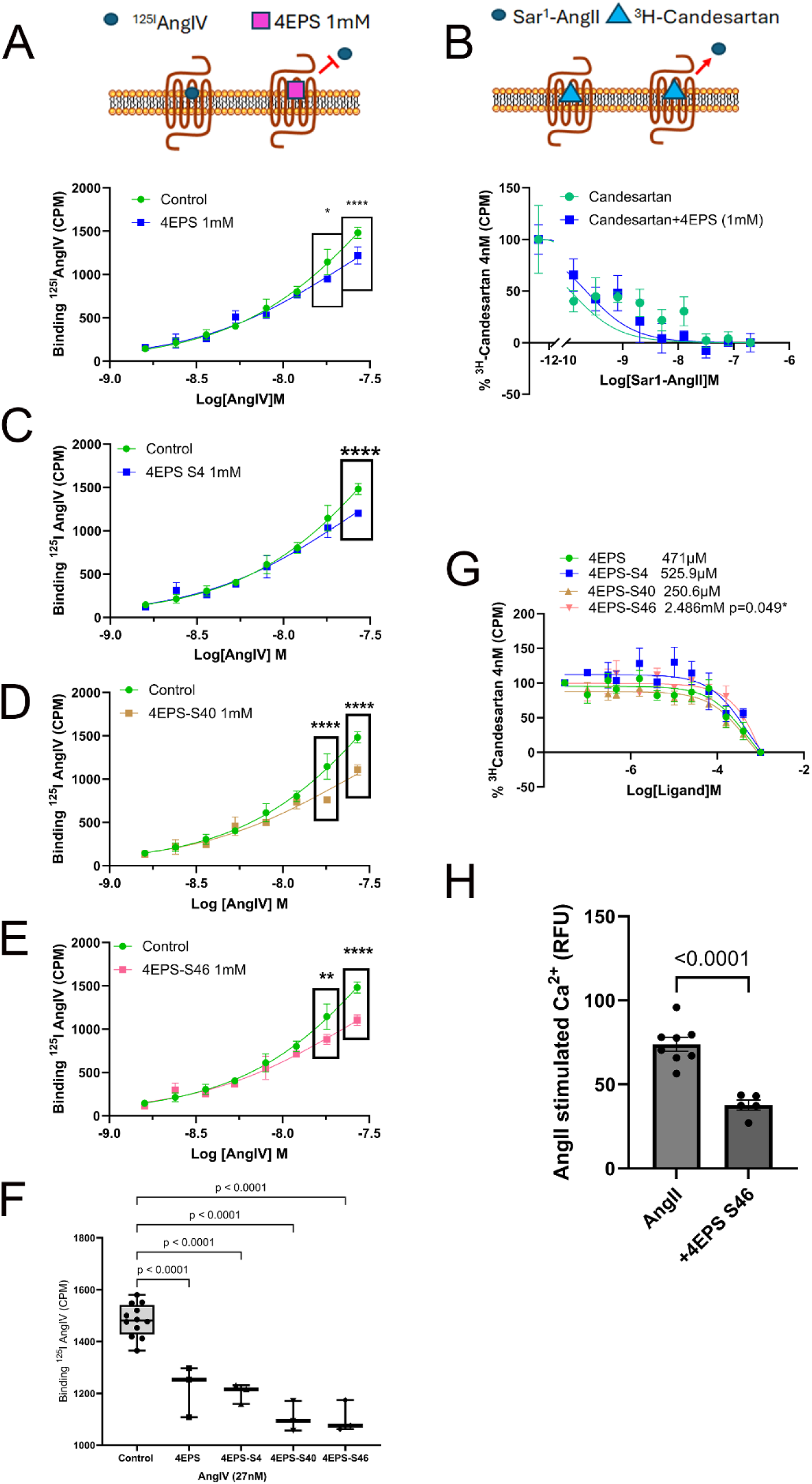
Effect of 4EPS and analogs on in vitro binding: Saturation binding assays were performed using membranes prepared from MOVAS-AT1R incubated with ^125^I-AngIV concentrations ranging between 1.6 and 27nM (specific activity, 16 Ci/mmol) with 1mM of A. 4EPS or C-E. 4EPS analogs (S4, S40 and S46) respectively. B Competition binding performed using membranes from HEK293-AT1R cells performed under equilibrium conditions using radioligand [^3^H]-Candesartan (4nM) and 0.04-1000nM concentrations of [Sar1] AngII with or without 4EPS (1mM). Two independent experiments, each done in triplicate, were performed and the resulting values were pooled to generate a mean curve which is displayed. The binding kinetics were analyzed by the nonlinear curve-fitting program GraphPad Prism 10. The means ± SEM for the LogIC_50_ values were calculated as described earlier. F. Analysis of significance of saturation binding using One way ANOVA with Dunnett test. Analysis was performed to determine significance at single doses in boxed data points in 2A,C-E which compare AngIV only and AngIV+4EPS or analogs using student t-test in GraphPad Prism 10. G. Competition binding performed as in 2B with 4EPS analogs in concentrations ranging from 1nM-1mM using radioligand [^3^H]-Candesartan (4nM). The binding kinetics were analyzed by the nonlinear curve-fitting program GraphPad Prism 10. The means ± SEM for the LogIC_50_ values were calculated as described earlier.

Molecular modeling and unbiased docking analyses indicated that 4EPS occupies a sub-pocket within the orthosteric ligand binding site of AT1R [**Fig 3** and **Fig S2**]. This sub-pocket accommodates the Phe^8^-COO^―^ sidechain of AngII and AngIV, as well as the biphenyl immidazole scaffold of angiotensin receptor blockers (ARBs), such as olmesartan [**Fig S2**]. Occupancy of this sub-pocket by 4EPS is facilitated by AT1R residues, Arg^167^, Lys^199^ and Trp^84^ [**Fig S2**]. Previous studies have shown that mutagenesis of Arg^167^, Lys^199^ and Trp^84^ produce non-functional AT1R protein, precluding mutagenesis approach to validate 4EPS-AT1R docking model. We developed an innovative pharmacological approach described below to circumvent this hurdle.

**Figure 3:**
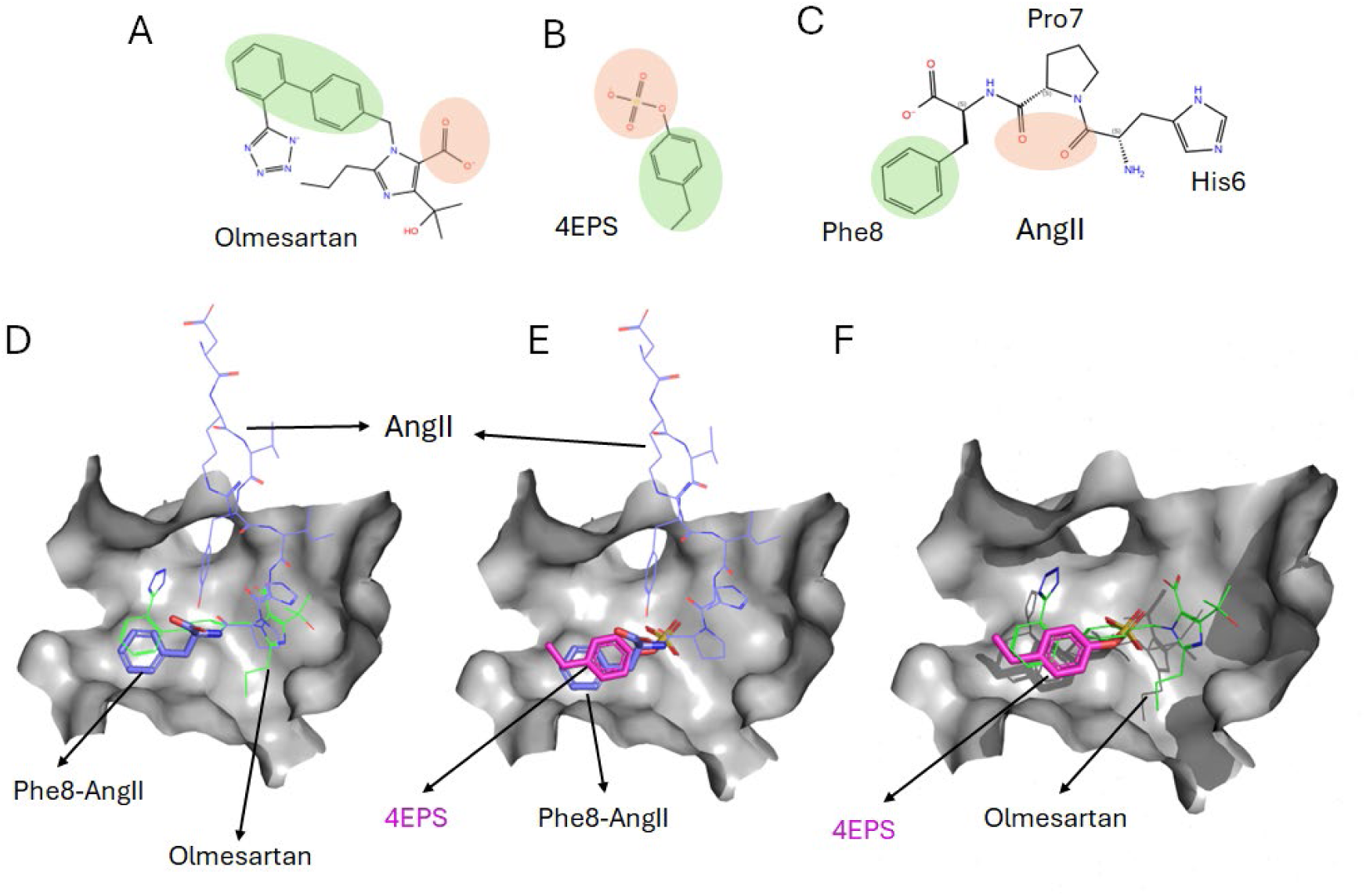
Model of 4EPS interaction with orthosteric pocket of AT1R. Chemical structure of Olmesartan (A), 4EPS (B), and the last three residues of AngII (C). Modeled docking poses of indicated ligands within the orthosteric ligand binding pocket of AT1R (D-F). The green regions represent hydrophobic areas, while the orange regions indicate hydrophilic areas. The ethylphenyl group of 4EPS occupies the position of Phe^8^ in AngII and the biphenyl position in Olmesartan. The sulfate group of 4EPS takes the place of the backbone carbonyls of Pro7-His6 in AngII and the carboxylic group of Olmesartan, indicating that 4EPS has inhibitory potential against both AngII and ARBs. The grey surface in D-F represents the protein inter-face of orthosteric pocket to which AngII, 4EPS and ARBs interact.

We searched chemical space using SciFinder, identified 54 4EPS-like compounds and performed Induced-Fit Docking (IFD). This analysis indicated that the docking score of five compounds [**Table S1**] was higher than 4EPS. Of these, only S4, S40, and S46 were commercially available as potential tools to pharmacologically interrogate the 4EPS binding site of AT1R. In ^125^I-AngIV binding analysis, S4, S40, and S46 (1mM) reduced the binding of radioligand [**Fig 2C-E**]. Order of inhibitory potency was S46 ≈ S40 > S4 ≈ 4EPS, which indicated that replacement of ethyl group in 4EPS by tertbutyl in S4 had minor effect and replacement with bulkier phenyl in S46 and benzyl in S40 had a larger effect [**Fig 2F**]. In the ^3^H-Candesartan tracer displacement, in the presence of 4EPS, S4, S40, and S46, inhibitory potency of S46 >S4≈4EPS>S40 was seen [**Fig 2G**], again showing S46 having the greatest effect in displacement. As compound S46 was most effective at displacement, we evaluated S46 effects on AT1R activation using the calcium assay and found that this compound containing benzyl substitution inhibited the response in HEK-AT1R cells.

### Effect of 4EPS on AngII-induced vascular reactivity *ex vivo*

Next, we evaluated 4EPS antagonism potential in a physiological system, aortic contraction. We measured *ex vivo* reactivity of mouse aortic explants from ApoE-null mice fed HFD and infused with different ligands as schematized in **Fig 4A**. Contractile responses were measured upon 300nM AngII-stimulation using wire myography. As described previously, integrity of endothelium and vessel-wall of aortic explants was assessed by measuring contraction/relaxation responses to treatment of high potassium physiological salt solution (KPSS) followed by phenylephrine and acetylcholine. Aorta from mice infused with AngII (1.4mg/kg/d) only produced a stronger contraction in response to AngII (300nM) stimulation [p=0.0016]. In contrast, mice infused with 4EPS (2.5mg/kg/d), AngII+4EPS and AngII+Olm elicit reduced contraction in response to AngII (300nM) stimulation [**Fig 4B**].

**Figure 4:**
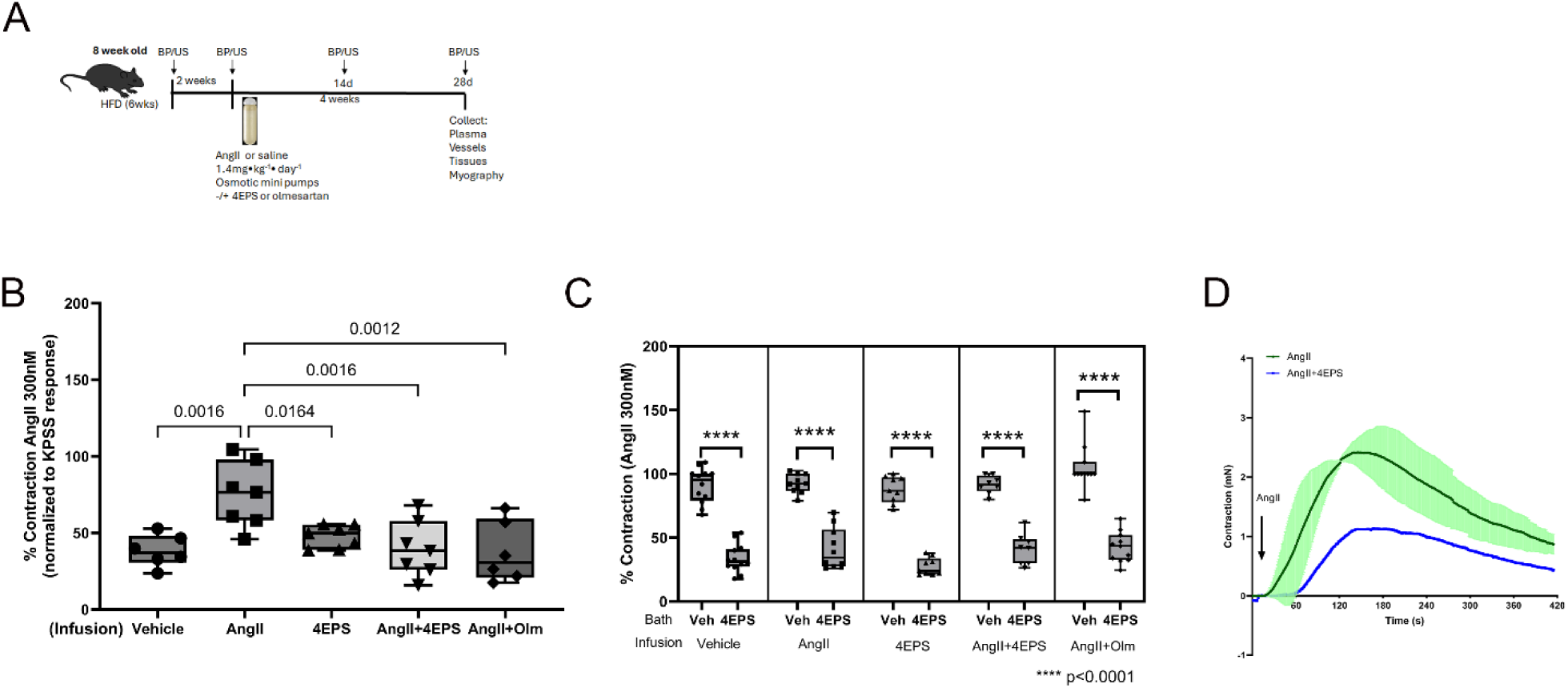
Ex vivo assessment of aortic contraction in response to AngII. A. Graphical experimental design. ApoE-null mice on HFD were infused with AngII (1.4mg/kg/d) or vehicle or 4EPS (2.5mg/kg/d) for 4 weeks. Wire myography was used to measure vascular contraction of infrarenal abdominal aorta in mice infused with ligands as indicated stimulated with AngII (300nM). Magnitude of contraction (mN, milli-Newton) (max-min) was normalized to maximal contraction stimulated by KPSS. Significance was calculated using One way ANOVA with Dunnett’s test comparing to vehicle infused using GraphPad Prism 10. C. Vasoconstriction was assessed with pretreatment with vehicle or 4EPS (100µM) prior to stimulation with AngII (300nM). Graph depicts percent contraction comparing constriction with AngII (300nM) pretreating with vehicle control or 4EPS. D. Graphical representation of contraction response in Labchart. AngII response (in green) is average of AngII response prior to pre-incubation with 4EPS and after AngII+4EPS response. Analysis of vehicle vs. pretreatment with 4EPS was performed using Student t-test.

We then determined the effect of acute 4EPS (100µM) addition in myography chamber on contraction. Preincubation with 4EPS (100µM) resulted in a significant reduction in AngII-mediated constriction regardless of the infusion [p=0.0001, in **Fig 4C**]. Both male and female ApoE-null mice fed a HFD were used for in the myography experiments, but female data is not shown due to poor response to AngII; for instance, the effect of AngII-stimulation was not significant in the aorta explants from female mice (p=0.4657). Graphical representation of contraction curves is depicted showing pre-incubation with 4EPS resulted in a reduction in AngII mediated vasoconstriction [**Fig 4D**]. Although aorta from both male and female mice were assessed, data for male is shown. Based on the lack of strong response in female subjects, we focused the remainder of our studies using male mice only.

### Effect of 4EPS on AngII-induced blood pressure

To determine in vivo effect of 4EPS, alterations in BP were measured by radiotelemetry monitoring. A transient elevation in mean arterial pressure in response to i.p. injection of AngII (0.2mg/kg) was observed in C57B6/J mice as compared to normal saline controls p=0.0096) [**Fig 5A**]. I.P. injection with a combination of AngII (0.2mg/kg) + 4EPS (0.4mg/kg) produced a transient increase in blood pressure not exceeding that of normal saline (p=0.9904). Area under the curve for each treatment group (n=3) indicates significant reduction of acute BP response [**Fig 5B**]. This experiment clearly shows the antagonist effect of 4EPS under physiological conditions. We next investigated the long-term effect of 4EPS utilizing ApoE-null mice fed HFD and infused over 28 days to initiate chronic pathophysiological conditions.

**Figure 5:**
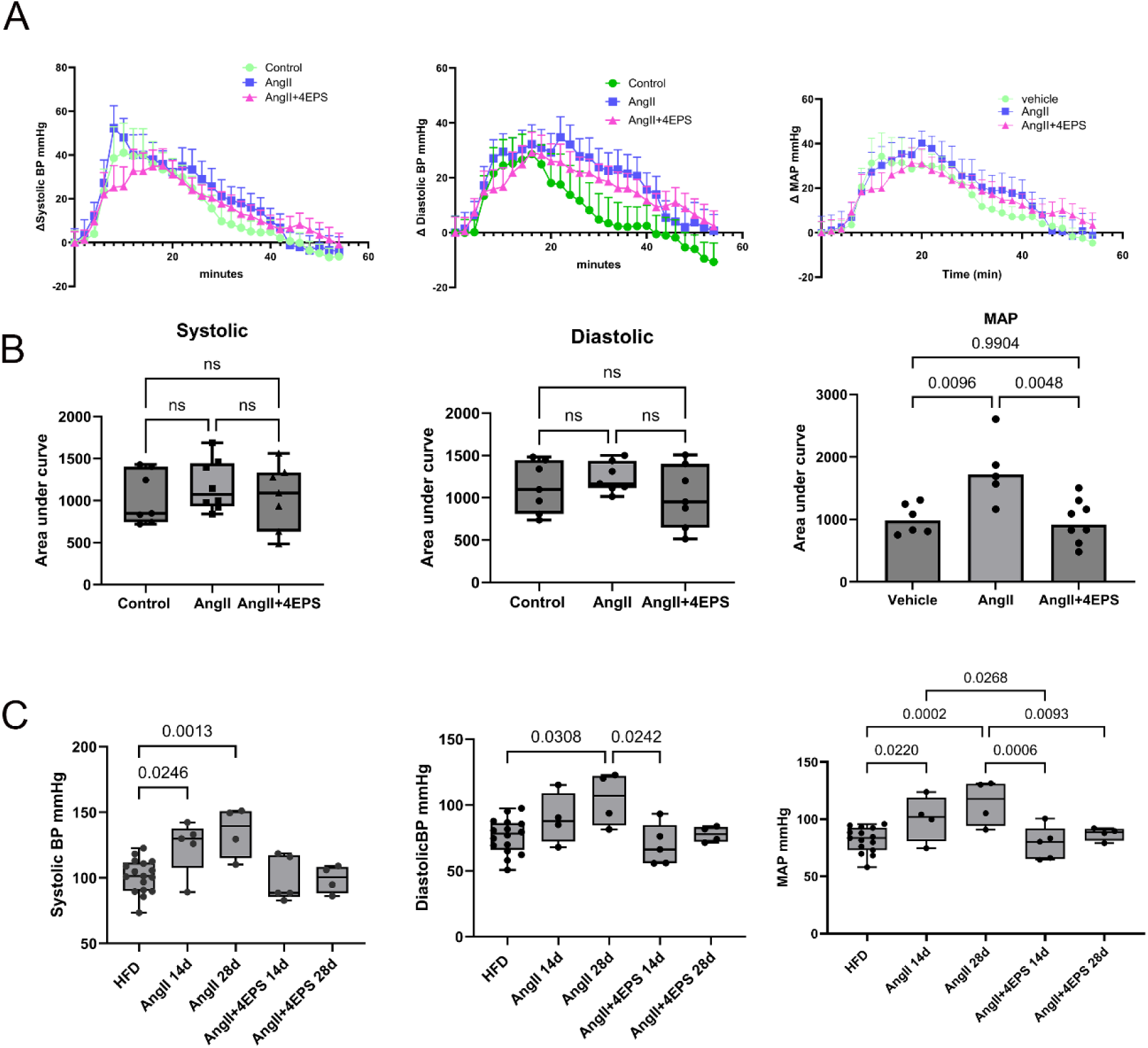
Diminished BP response in vivo. A. Systolic, diastolic and mean arterial pressure monitored in response to acute treatment. C57B6 injected with normal saline, AngII (0.2mg/kg), AngII+4EPS (0.2mg/kg and 0.4mg/kg respectively) while pressure was monitored by implanted radiotelemetry devices in conscious freely moving mice 20 min prior to and 60 min post injection. 3 mice per group with triplicate treatments over 5-day period to allow compound washout. B. Area under curve analysis was performed using Graphpad Prism 10 using One way ANOVA to determine significance. C. Alterations to blood pressure was also measured in response to chronic exposure in ApoE-null male mice. Mice on HFD were infused with vehicle (saline), AngII (1.4mg/kg/day), AngII+4EPS (AngII 1.4mg/kg/day, 4EPS 2.5mg/kg/day) 4 weeks. Pressure was measured during HFD followed by 14d and 28d post infusion by CODA non-invasive tail cuff. Analysis performed by One Way ANOVA with Tukey post-test using GraphPad Prism 10

Next, we measured BP changes in mice using non-invasive tail cuff monitoring using the chronic infusion protocol as described in **Fig 4A**. BP readings were taken 10 days into HFD, 14 days after osmotic pump implantation and finally at 28 days post implant. Systolic blood pressure significantly increased at 14 (p=0.0246) and 28 days (p=0.0013) of AngII infusion, diastolic pressure increased at 28 days (p=0.0308) and mean pressure increased significantly at 14 (p=0.0220) and at 28 days (p=0.0002) AngII infusion when compared to 10 days of HFD. In mice co-infused with AngII+4EPS, there was no significant increase in all three BP parameters [**Fig 5C**]. Together, these observations suggest that co-infusion with 4EPS blocked the hemodynamic effects of AngII similar to antagonist effect of the ARB, Olmesartan.

### Effect of 4EPS on AngII-induced aortic aneurysm in ApoE-null mice

We determined the effects of 4EPS on AngII-induced AA in ApoE-null male mice maintained on HFD. Five 28-day treatment groups described in methods received either vehicle or AngII, AngII+4EPS, 4EPS alone or Olmesartan. 100% mice survived in the vehicle (n=12) and 4EPS alone (n=7) and Olmesartan (n=6) groups. In the AngII group (n=16), 54% mortality was recorded. The mortality in the AngII+4EPS group (n=14) was 18%, suggesting substantial (p=0.0488) protection by 4EPS (**Fig 6A**). Monitoring weights of liver, spleen, lungs, heart and kidneys normalized to overall body weight indicated that co-infusion with 4EPS did not cause any change. However, overall body weight was reduced in AngII, AngII+4EPS and 4EPS only groups; the body weight alterations may be due to lean/fat mass changes (**Fig S3**). In the female ApoE-null mice infused with AngII, co-infused with 4EPS or ARB no significant change was observed.

**Figure 6:**
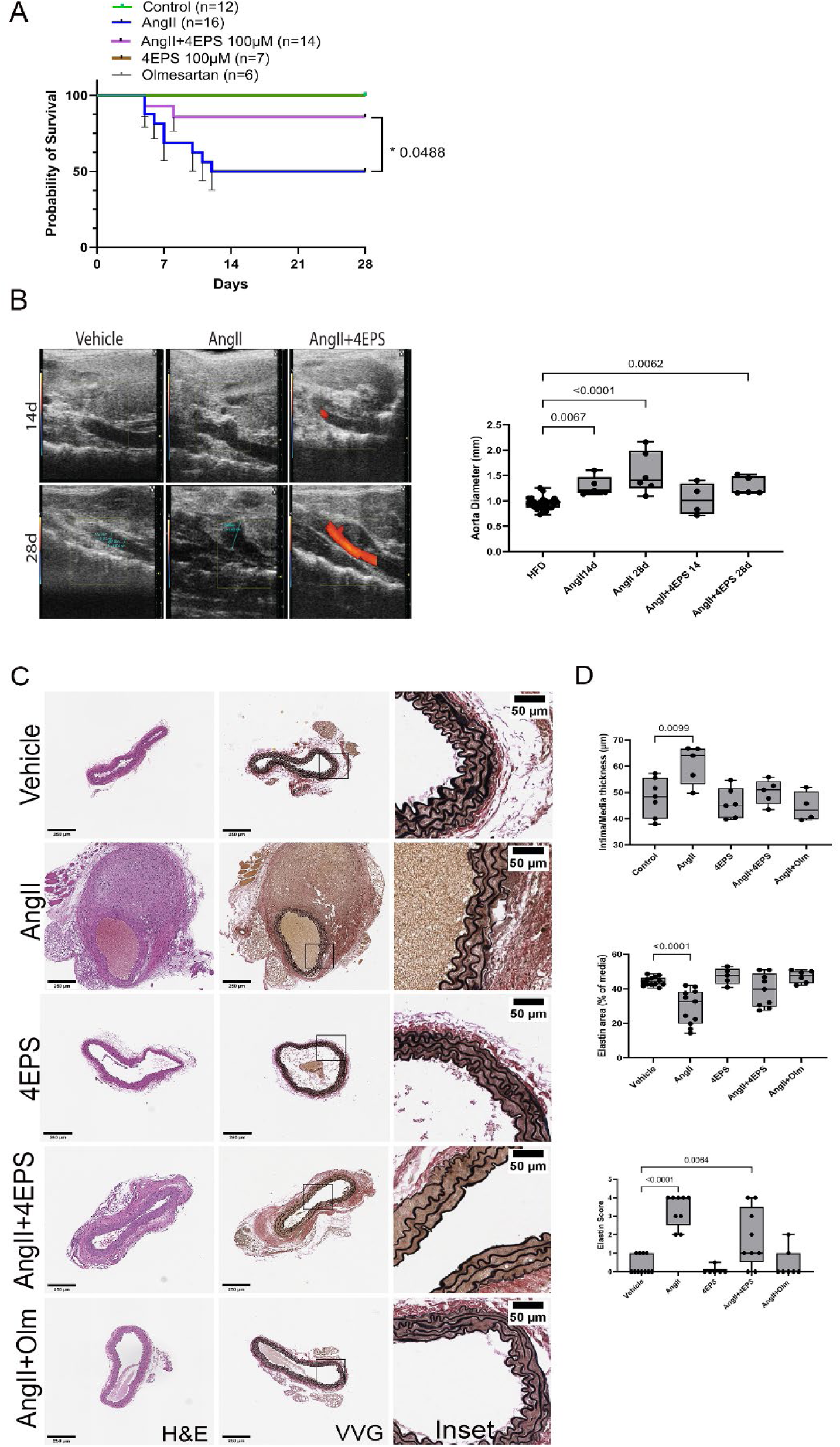
Infusion of 4EPS reduces AngII induced aneurysm and increased survival. ApoE-null mice infused as indicated for 28 days while on HFD. A. Kaplan-Meier simple survival curves from ApoE-null mice infused as indicated were generated in GraphPad Prism 10 using Log rank test for significance n=6-16 based on treatment). B. Representative long axis ultrasound images of aortic lumen diameter was monitored at same times as BP. Images show aorta with lumen outlined in red. Diameters were measured using Vevo Lab software on images acquired on Vevo2100 using ms550 probe. Significance was determined using One way ANOVA and Tukey multiple comparison test using GraphPad Prism 10. Histological analysis of Aorta intima/media thickness was measured H&E-stained tissues. Representative histological images of tissues. Scale bar 300µm. Elastin content was measured, and scoring was performed on Verhoef van Gieson stained tissues and scoring was performed by 4 independent reviewers of deidentified images. Analysis was performed using One way ANOVA with Dunnett’s multiple comparison test using GraphPad Prism 10.

AA growth progression in mice was monitored by imaging the maximal aortic lumen diameter using ultrasonography using Vevo2100 echocardiography system (Visualsonics). AngII infusion led to increased suprarenal abdominal aortic diameter at both 14-days (1.5-fold increase, p=0.0133) and 28-days (2-fold increase, p<0.0005). In contrast, significant increase (1.3X increase, p=0.0147) in lumen diameter was observed at only 28-days which suggests delays in progression of abdominal aortic aneurysms in the AngII+4EPS group (**Fig 6B**). Histopathology analysis of suprarenal abdominal aortic sections demonstrated a significant (p=0.0099) thickening of the aortic wall intima/media layers associated with luminal thrombus in AngII infused mice (**Fig 6C**). We assessed evidence of elastolysis by monitoring changes in elastin content and structural alterations within the tunica media of the aorta. Elastin content measurement showed a significant decrease in the AngII infused group compared to AngII+4EPS, 4EPS, AngII+ Olm and the vehicle control groups (p<0.0001). Infusion with 4EPS, and AngII+4EPS and AngII+Olm did not significantly differ as compared to vehicle control group (p=0.9711, p=0.9996, p=0.8832 respectively) (**Fig 6D**).

### Effect of 4EPS on molecular pathways underlying AngII-induced aortic aneurysm

Unbiased proteomic analysis on the plasma of ligand infused mice identified potential molecular signaling mechanisms responsible for reduced vascular remodeling in the AngII+4EPS compared to AngII treatment groups. Significantly altered proteins are shown in Table S2. The proteomic data was analyzed by IPA, String and GO to identify altered canonical pathways, diseases and biological functions in response to our treatments. Changes identified to key canonical pathways were in actin cytoskeleton signaling, and ERK/MAPK signaling. Alterations in biofunction pathways involving changes in chemotaxis of blood cells (granulocyte, neutrophils, phagocytes) adhesion property of vascular endothelial cells, production of NO and ROS were observed. In the String database analysis, the top hit identified was molecules associated with elastin fibers and kidney disease. GO analysis elucidated changes associated with integrin signaling, blood coagulation and inflammation [**Fig S4**].

The predicted 4EPS influence on AngII-induced actin cytoskeletal signaling may be relevant to understand vascular wall remodeling changes observed in the 4EPS+AngII co-infused mice in **Fig 6**. Using the MOVAS-AT1R cells, we monitored cell-migration capability (see methods) in a wound-healing assay shown in **Fig S5**. MOVAS-AT1R cells, cultured in serum free media, were incubated with saline control, AngII (1µm) or AngII+4EPS (100µm) over 24 hours. Images were acquired at 0h, 8h and 24h. Cell migration is observed in all treatment conditions. However, AngII+4EPS treatment diminished migration of cells when compared to the AngII alone treatment which substantially increased migration capability in comparison to saline control.

Next, we monitored alterations in expression and phosphorylation of Filamin A, a key actin cross-linking biomechanical signal transducer protein, which has been associated with changes in hypertension and CVDs^49,50^. Agonist activation increases Filamin A phosphorylation and recruitment to AT1R, which is known to be crucial for regulating the dynamics of actin cytoskeleton. MOVAS-AT1R cells were treated with AngII or AngII+4EPS and assessed for FlnA expression and phosphorylation [**Fig S6A**]. AngII treatment increased phosphorylation of Filamin A in comparison to treatment with AngII+4EPS. There was no significant effect on Filamin A expression. These data suggest that 4EPS directly inhibits AT1R− Filamin A-cytoskeleton coupling induced by AngII. As the proteomic analysis suggested alterations in ERK/MAP kinase signaling, ERK expression and phosphorylation was assessed [**Figure S6B and C**]. Again, agonist activation transiently increases ERK phosphorylation in response to activation of AT1R which is critical for the transient increase in Ca^2+^_i_, that stimulates smooth muscle contractions. MOVAS AT1R and HEK-AT1R cells were treated with AngII or AngII+4EPS. In both cell lines AngII treatment led to an increase in phosphorylation of ERK as compared to AngII+4EPS.

Our plasma proteomics data indicated an inverse association of 4EPS with chronic renal impairment, in contrast to literature reports that indicated positive correlation with chronic kidney disease and end stage renal disease. We assessed circulating 4EPS levels in our experimental mice by measuring plasma 4EPS by mass spectrometry. Interestingly, plasma 4EPS levels were significantly higher in AngII+4EPS co-infused mice (p<0.0001) than 4EPS only infused mice. As anticipated, the 4EPS level was below the level of detection in vehicle control and in AngII-infused mice. This observation led us to speculate the possibility that renal clearance of 4EPS is inhibited in AngII+4EPS co-infused mice thereby indicating potential kidney dysfunction. To evaluate potential renal dysfunction, we assessed markers of kidney damage by monitoring mRNA levels of Col1A1, TGFb, KIM-1 and SMA [**Fig S7**]. As compared to vehicle infused mice, there was no change in Col1A1 or TGFβ between AngII, AngII+4EPS and AngII+Olmesartan, however AngII infusion led to a significant increase in KIM-1 and SMA, with no concurrent change in AngII+4EPS infusion. These observations indicate retention of 4EPS in circulation in the AngII+4EPS co-infused mice but lack of 4EPS associated renal injury.

## Discussion

Our lead-off question in this study was whether the microbial metabolites reported in the Cheema and Pluznick study^19^ modulate AT1R signals directly responsible for vaso-regulatory actions of AngII? We used the HEK-AT1R cell-signaling platform, in which AngII stimulation of cells elicits a transient and saturable increase in cytosolic calcium. In this assay, addition of metabolites is expected to alter the EC_50_ and/or the E_max_ of the AngII dose-response curve. Six microbial metabolites elevated upon AngII infusion in normal mice compared to germ free mice were screened [**Fig 1**]. Only AngII+4EPS combination led to a significant EC_50_ reduction compared to the AngII only control measuring AT1R activation. Lack of a significant inhibitory effect by the other five metabolites suggested that 4EPS inhibition of AT1R signaling is specific. Furthermore, the calcium response to both AngII and AngII+4EPS were dependent on AT1R expression in HEK cells. The AT1R selective drug, Olmesartan inhibited more effectively when compared to inhibition by 4EPS. Thus, 4EPS acts as a “benign antagonist” of AngII signaling. It is known that the production of 4EPS from catabolism of phenylalanine and tyrosine is regulated by gut microbial metabolism of dietary protein and aromatic amino acids^10^. Elevation of plasma 4EPS levels linked to type 2 diabetes mellites^8^, chronic heart failure^11^, neurological changes^40^ and chronic kidney diseases^38^ is reported in humans and mice. However, the impact of 4EPS on the RAS and particularly its molecular target is not known. Thus, demonstrating “benign antagonism” of AT1R by 4EPS is a significant advance we report here.

AT1R induced calcium generation is a second/third messenger signaling system of hormonal actions of AngII. It’s inhibition by 4EPS confirms its functional interaction with AT1R, but not whether 4EPS binds to AT1R. We examined this question by in vitro ligand binding studies. Specific binding of the partial agonist, ^125^I-AngIV to membranes isolated from MOVAS-AT1R cells is systematically reduced by 4EPS [**Fig 2A**]. In ARB vs AngII competitive binding experiment, displacement of the ARB, ^3^H-Candesartan by increasing concentrations of [Sar1]AngII was accelerated (left-shifted the curve) in the presence of 4EPS [**Fig 2B**]. This suggests that 4EPS interferes with the binding of both ARB and AngII. Previous structural and mutagenesis studies established well defined binding sites of ARBs and AngII in AT1R ^42,51,52^. Our unbiased in silico docking of 4EPS to crystal structure of AT1R indicated that 4EPS binding to AT1R involved specific AT1R-residues, Arg^167^, Lys^199^ and Trp^84^ within the orthosteric binding pocket of AT1R [**Fig 3, Fig S2B**].

These residues interact with all orthosteric ligands which explains the basis of 4EPS inhibition of binding of AngIV, AngII and Candesartan. Furthermore, the docking analysis showed that 4EPS occupies the same space that accommodates Phe^8^ residue of AngII within the orthosteric pocket of AT1R. It is well established that Phe^8^ in AngII is most critical for activation of AT1R^53^, and 4EPS inhibits AT1R activation by hindering Phe^8^ interaction with AT1R.

By employing 4EPS-analogs [**Fig S2A**], we established further that 4EPS occupies a confined sub-pocket responsible for accommodating Phe^8^ of angiotensin. 4EPS-analogs with bulkier benzyl and phenyl substitutions inhibited specific binding of ^125^I-AngIV more strongly than those with smaller ethyl and tert-butyl substitution [**Fig 2C-D**]. Increase in the inhibitory potential of 4EPS-analogs is consistent with stronger binding predicted by their docking scores. Taken together, our studies establish that 4EPS directly binds to the orthosteric pocket of AT1R and attenuates AngII-induced calcium signaling and perhaps additional downstream effects.

We next investigated potential physiological effects of 4EPS, by measuring vascular response utilizing ex vivo wire myography [**Fig 4B**]. The significantly higher magnitude of contraction (p=0.0016) produced in the AngII infused mice was diminished by 4EPS. Pre-incubating aortic explants with 4EPS also showed significantly reduced contractile response to AngII, ∼50% reduction (p<0.0001). Inhibition of contraction of aortic explants by 4EPS is consistent with inhibition of calcium response in cell culture suggesting potential humoral impact of 4EPS on host physiology.

As this is the first study to determine the effect of 4EPS on cardiovascular health, we investigated its *in vivo* effects on AngII-induced AA development. A commonly used model for experimental AA studies, depends on AngII-mediated activation of AT1R in the context of ApoE-null genotype and high-fat diet to generate the AA pathophysiological state that mimics clinical findings that hypercholesterolemia, obesity, and hypertension increase the risk of AA mortality ^54–56^. Therefore, 4EPS was tested in this model for efficacy to prevent the disease. While the systolic BP was elevated in response to 14- and 28-days of AngII infusion, there was no concurrent rise of BP in AngII+4EPS co-infused mice. Both diastolic and mean BP was elevated in AngII group at 28 days with no significant changes in AngII+4EPS co-infusion group [**Fig 5**]. Mass spectrometry analysis ascertained that the plasma levels of 4EPS increased significantly in the 4EPS and AngII+4EPS infused mice [**Fig S7A**], which supports the conclusion that attenuation of BP increase by AngII is associated with increased plasma levels of 4EPS following osmotic-pump infusion. Thus, 4EPS prevented AngII-induced elevation of BP validating the results of the ex vivo myography measurements.

Prevention of BP elevation upon 4EPS co-infusion in ApoE-null mice was associated with inhibition of AngII-induced growth of AA and structural alterations within abdominal aortic wall [**Fig 6**]. Thickening of the intima/medial layer of abdominal aorta is indicative of vascular remodeling and has been identified as a precursor to the development of AA^56^. Histological comparison of aortas of AngII+4EPS co-infusion group identified significant prevention of thickening of the intima/media layers compared to that observed in the AngII infused group (p=0.0099). The vessel wall elastin content in the AngII+ 4EPS co-infused mice resembled vehicle treated mice, while AngII infusion led to a significant reduction in elastin content (p=0.001). Reduced elastin content reflects elastolysis and weakening of aortic wall which has been associated with increased aneurysm dissection, hypertension and congenital disorders including Marfan Syndrome^57–59^ Ultrasound imaging analysis showed a significant increase in aortic diameter in 14-days and progressed until 28-days in the AngII infused group, whereas in the AngII+4EPS co-infusion group slight increase in diameter was observed at 28-days post infusion. No significant changes were observed in organ weight (normalized to body weight) in liver, spleen, lung, kidneys and heart. Aortic wall remodeling data lead us to speculate that AngII+4EPS co-infusion delays AA formation, which is also reflected in overall changes in mortality. We have previously shown that AngII infusion induces AA located in the suprarenal abdominal aortic region in 64% male ApoE-null mouse fed high fat western diet^41^. In this study, we found abdominal AA death in 53% mice when infused with AngII, which was reduced to 14% when co-infused with 4EPS. The overall survival is significantly improved in mice co-infused with AngII+4EPS as compared to AngII alone (p=0.0488).

Molecular mechanistic insight regarding inhibitory effect of AngII+4EPS co-infusion on AA growth is provided by unbiased proteomic analysis of plasma proteins [**Table S2**]. The altered molecular changes revealed that 4EPS treatment attenuated AngII-induced actin cytoskeletal signaling, which is relevant to vascular wall preservation observed in the 4EPS+AngII co-infused mice. We directly tested migration of cells and observed that AngII alone treatment substantially increased migration capability of cells and AngII+4EPS treatment reduced cell migration [**Fig S5**]. In addition, Filamin A phosphorylation [**Fig S6**] was increased by AngII treatment in comparison to treatment with AngII+4EPS. Filamin A is a key actin cross-linking and biomechanical signal transducer protein. In a previous study^49^, we demonstrated that direct high-affinity interaction of Filamin A with the filamin binding motif of AT1R located on its helix-8 regulates biomechanical signaling by AT1R. Engagement of filamin binding motif of AT1R with Filamin A induces the phosphorylation of filamin in cells. In the current study AngII increased the phosphorylation of cellular Filamin A, that is crucial for regulating cytoskeleton dynamics and enhanced cell migration. Recent genetic and molecular studies have shown a direct role of Filamin A in aortic aneurysm pathology in humans and mice^50^. By interfering with AT1R activation by AngII, 4EPS might attenuate biomechanical signaling by AT1R to actin-cytoskeleton of aortic wall cells and prevent remodeling.

Experimental demonstration that AT1R is the molecular target of 4EPS provides basis for mechanistic link to RAS and cardiovascular diseases. It is one of a dozen possible gut microbial metabolites produced by the participation of microbial and host enzymatic pathways. For instance, in producing 4EPS from dietary protein, 4-hydroxy phenylpropionic acid is produced by gut-microbes, followed by sulfonation reaction carried out by the host liver enzymes. Clinical studies have identified several metabolites associated CVD risks, molecular targets for most of these metabolites are not established. In this context our demonstration that AT1R as the target of 4EPS is highly significant for considering risk benefits evaluation for cardiovascular diseases. 4EPS link to neurological changes may also involve AT1R ^40,60–63^. By crossing the blood-brain barrier 4EPS interaction with AT1R may influence neural function and brain activity for example related to Autism spectrum. Studies have demonstrated link between microbial alterations in several neurological and psychiatric disorders, including anxiety, depression, and neurodegenerative diseases ^40,61,63^. Moreover, 4EPS may now be recognized as a component of the gut-brain axis implicated in the regulation of stress responses and the modulation of the immune system ^64^.

We noted that 4EPS link to chronic kidney diseases observed in humans and mice is of interest for future studies. In our experiments, we observed a 500% increase in 4EPS levels in the AngII+4EPS co-infused mice when compared to 4EPS infusion alone [**Fig S7**]. However, analysis of kidney injury markers suggested that infusion with AngII leads to kidney dysfunction and co-infusion with 4EPS attenuated this effect in mice. Further studies are needed to clarify mechanism of 4EPS attenuation of AngII-mediated kidney damage.

In summary, our in vitro, ex vivo and in vivo data together support the conclusion that AT1R is the molecular target of 4EPS. Both the calcium messenger signaling and the biomechanical signal transduction through protein-protein interaction triggered by AT1R are attenuated by 4EPS.

## PERSPECTIVES

Hypertension is associated with vascular diseases, kidney dysfunction and aneurysm formation. Gut bacterially derived metabolites have been shown to play a role in CVD. The metabolite 4EPS, has not been linked to hypertension or implicated in the regulation of RAS, which is the critical neurohormonal pathway needed to regulate hypertension. Our results demonstrate that 4EPS can attenuate AT1R activation by AngII, thereby reducing AngII mediated hypertension in male mice. When male mice are treated with 4EPS with AngII, blood pressure is reduced, aneurysm formation slows, and overall survival from AA is increased.

Our observation that 4EPS interferes with the binding of ARB implies that plasma 4EPS levels may be an influencing factor for ARB therapy application clinically.

Manipulating gut microbial functional output to produce beneficial metabolites like 4EPS can be a novel approach for consideration of potential therapeutic approaches treating CVD in future.

## Novelty and Significance

### What is new?

4EPS can selectively bind to AT1R and reduce AngII binding within the orthosteric pocket.

4EPS can reduce vascular contraction in response to AngII ex vivo.

Demonstrates for the first time that 4EPS can attenuate elevated blood pressure in response to AngII infusion in male mice.

Co-infusion with 4EPS and AngII can attenuate aortic abdominal aneurysm formation and associated mortality.

### What is relevant

The results of this study improve our understanding of the impact of bacterial derived metabolites, specifically 4EPS, on cardiovascular health, hypertension and aneurysm formation.

### Clinical/pathophysiological significance

Moreover, this study has clinical/pathophysiological significance in that hypertension can be affected by 4EPS thereby slowing aneurysm formation and kidney damage associated with hypertension because of increased RAS activity.

This study has translational significance in that 4EPS, and derivatives can used to design new AT1R inhibitors to better control hypertension in men.

## Supporting information

Manuscript text and Figures

## Nonstandard Abbreviations and Acronyms

4EPS: 4-ethylphenylsulfate
ACE: Angiotensin converting enzyme
Acta2: Alpha actin 2
AngII: angiotensin II (Asp-Arg-Val-Tyr-Ile-His-Pro-Phe)
AngIV: angiotensin 3-8 (Val-Tyr-Ile-His-Pro-Phe)
AT1R: angiotensin II type 1 receptor
OLM: AT1R inverse agonist, olmesartan
ARB: angiotensin II type 1 receptor (AT1R) blocker
AA: aortic aneurysm
AAD: aortic aneurysm dissection
ACEi: angiotensin converting enzyme inhibitor
ApoE: apolipoprotein E gene.
BP: blood pressure
Cola1a: Collagen a1a
Col3A1: Collagen 3a1
CVD: Cardiovascular Disease
eGFR: estimated glomerular filtration rate
ERK: Extracellular regulated kinase
FlnA: Filamin A
GPCR: G-protein coupled receptors
H&E: Hematoxylin & Eosin staining
HFD: High fat diet
IFD: Induced-Fit Docking
IPA: Ingenuity Pathway analysis
KIM-1: Kidney injury marker 1
KPSS: high potassium physiological salt solution
LFQ: label-free quantitation
MAPK: Mitogen activated protein kinase
MOVAS: Mouse aortic vascular smooth muscle
MT: Masson’s trichrome staining
RAS: Renin angiotensin system
SCFA: Short-chain fatty acids
SMA: Smooth muscle actin
TGFb: Transforming growth factor beta
VVG: Verhoeff-vanGieson staining
VSMC: vascular smooth muscle cells

## Acknowledgements

The authors would like to express their appreciation to the Proteomics core for their assistance in LC-MS analysis and the Imaging Core for their assistance with analysis of histological images. We thank Renliang Zhang for assistance with LC-MS quantitation of 4-EPS, and Ling Li for assistance with the quantitative proteomics. We thank the LRI core facilities including the Animal Core Facility, the Histopathology and Biomedical Engineering Imaging Core. We are especially indebted to Maya Camhi for immunohistochemistry assistance and Judith Drazba, for assistance in imaging and image analysis studies.

## Supplemental Material

Tables S1 and S2

Figures S1-S7

Graphic Abstract

