## Supplementary material for "Microbial metabolite 4-ethylphenylsulfate (4EPS) interacts with AT1R, reduces blood pressure and outcome of AngII-induced aortic aneurysm": Manuscript text and Figures

Terri J. Harford, Dhanachandra Khuraijam Singh, Triveni R. Pardhi, Russell Desnoyer, Tarun Ravi, Zaira Palomino Jara, Ajay Zalavadia, Kate Stenson, Sathyamangla V Naga Prasad, Sadashiva S. Karnik*

Cardiovascular and Metabolic Sciences Department, Lerner Research Institute, Cleveland Clinic

* Address Correspondence to:

Sadashiva S. Karnik, Ph.D.

Cardiovascular and Metabolic Sciences, Room NB50-76

Cleveland Clinic, Cleveland, OH 44095, USA

**Table S1**

| Code | IFD docking score [kcal/mol] |
|---|---|
| 4EPS | -6.97 |
| 4EPS_S4 | -7.839 |
| 4EPS_S19 | -8.427 |
| 4EPS_S37 | -8.167 |
| 4EPS_S40 | -7.942 |
| 4EPS_S46 | -8.616 |

**Table S1:** First performed XP docking of all 54 compounds followed by Induced fit docking (IFD) of all 54 compounds. Identified 5 compounds, S4, S19, S37, S40, and S46 with docking score >10-fold the docking score of 4EPS.

**Table S2**

| Protein | Accession | Gene ID | LFQ ratios | |
|---|---|---|---|---|
| | | | AngII+4EPS/AngII | p-value |
| 60S ribosomal protein L15 | Q9CZM2 | Rpl15 | 5.160 | 0.052 |
| ATP-dependent 6-phosphofructokinase, liver type | P12382 | Pfkl | 3.493 | 0.059 |
| Ubiquitin thioesterase OTUB1 | Q7TQI3 | Otub1 | 2.529 | 0.043 |
| Beta-1,4-galactosyltransferase 1 | P15535 | B4galt1 | 2.508 | 0.057 |
| Ig kappa chain V-V region K2 (Fragment) | P01635 | | 1.894 | 0.045 |
| Fatty acid-binding protein, adipocyte | P04117 | Fabp4 | 1.880 | 0.014 |
| Ig kappa chain V-III region 50S10.1 | P03977 | | 1.800 | 0.014 |
| Microfibril-associated glycoprotein 4 | Q9D1H9 | Mfap4 | 1.671 | 0.028 |
| Glutathione S-transferase omega-1 | O09131 | Gsto1 | 1.634 | 0.055 |
| MLV-related proviral Env polyprotein | P10404 | | 1.560 | 0.023 |
| Ceruloplasmin | Q61147 | Cp | 1.197 | 0.001 |
| Vitronectin | P29788 | Vtn | 0.897 | 0.039 |
| Retinoic acid receptor responder protein 2 | Q9DD06 | Rarres2 | 0.896 | 0.041 |
| Hepatocyte growth factor activator | Q9R098 | Hgfac | 0.820 | 0.036 |
| Alpha-1-antitrypsin 1-1 | P07758 | Serpina1a | 0.809 | 0.015 |
| Angiotensin-converting enzyme | P09470 | Ace | 0.791 | 0.018 |
| Serine protease inhibitor A3C | P29621 | Serpina3c | 0.788 | 0.018 |
| Tetranectin | P43025 | Clec3b | 0.778 | 0.002 |
| Interleukin-1 receptor accessory protein | Q61730 | Il1rap | 0.757 | 0.039 |
| 72 kDa type IV collagenase | P33434 | Mmp2 | 0.728 | 0.021 |
| Rab GDP dissociation inhibitor alpha | P50396 | Gdi1 | 0.687 | 0.047 |
| Carboxylesterase 3A | Q63880 | Ces3a | 0.687 | 0.044 |
| Ubiquitin carboxyl-terminal hydrolase 5 | P56399 | Usp5 | 0.622 | 0.008 |
| Integrin beta-1 | P09055 | Itgb1 | 0.510 | 0.056 |
| Odorant-binding protein 1a | Q9D3H2 | Obp1a | 0.343 | 0.027 |
| Triokinase/FMN cyclase | Q8VC30 | Tkfc | 0.086 | 0.003 |

**Table S2: Proteomic analysis of plasma proteins in 4EPS+AngII vs AngII infused mice** (n=3). Quantitative Unbiased proteomic analysis of plasma of AngII Vs Vehicle and 4EPS+AngII vs Vehicle was performed. Protein quantities were expressed as normalized intensities by Spectronaut V17. Summary 21 proteins exhibited significant changes between AngII vs AngII+4EPS, 7 increased with AngII and 14 decreased with AngII infusion. LFQ -label free quantitation.

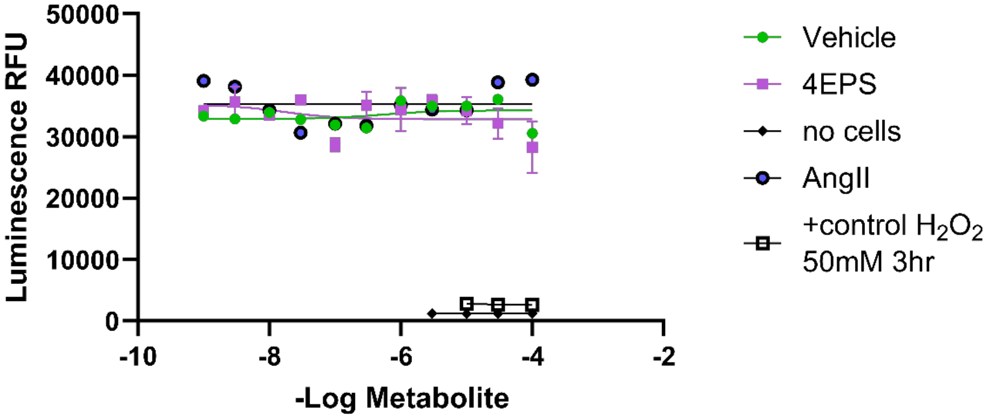


**Figure S1: 4EPS exhibits no cytotoxic effects.** To ensure this effect was not due to cytotoxicity, MOVAS cells were incubated for 24 hours in the presence of vehicle, 4EPS or AngII (doses 1nM-100μM) or $H_2O_2$ (50mM 3hrs, positive control) no cells used as blank. Cell viability was measured using CellTiter Blue Cell viability assay. Data represents mean± SD. N=2 samples run in duplicate

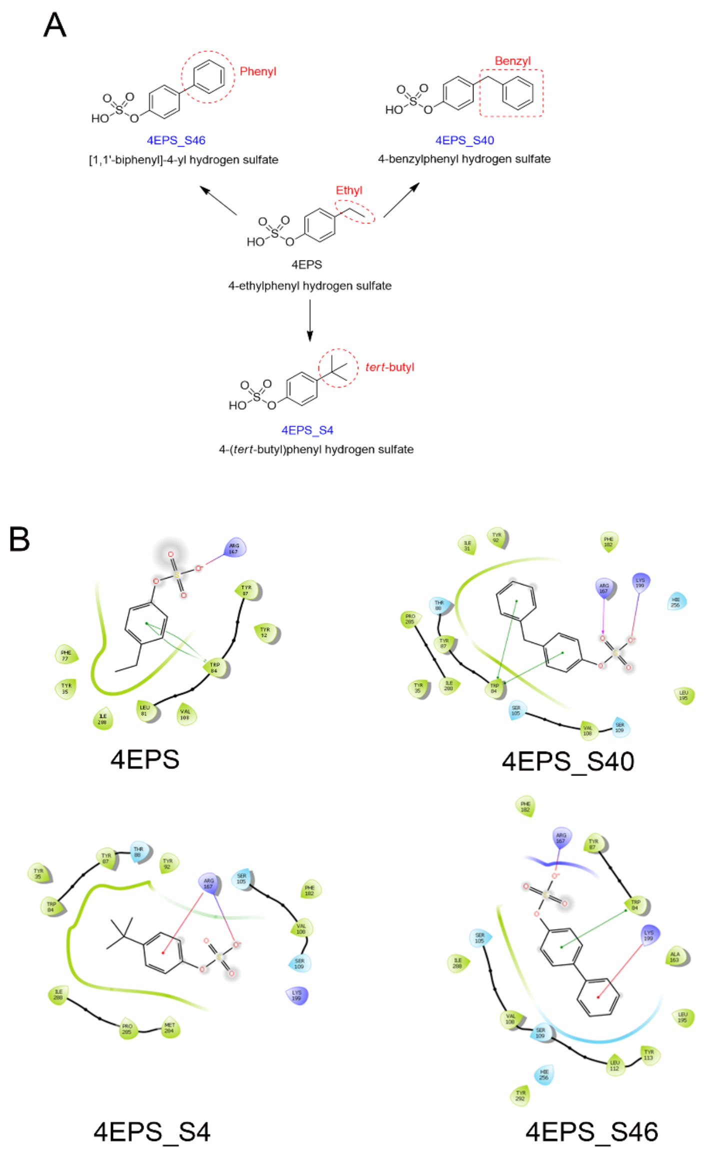


**Figure S2: A. Structural relationship of 4EPS analogs.** Ethyl group at 4 positions in 4EPS is substituted with phenyl, benzyl, and *tert*-butyl in S46, S40, and S4 respectively. We selected similar compounds and assessed alterations in AT1R activation by changes in cytosolic calcium levels. B. The docking interactions of these compounds utilize residues $Arg^{167}$, $Lys^{199}$ and $Trp^{84}$ for binding in the orthosteric ligand pocket of AT1R, which are exactly the residues that are essential for binding the ARB Losartan as shown and other sartan drugs as shown by us previously[49].

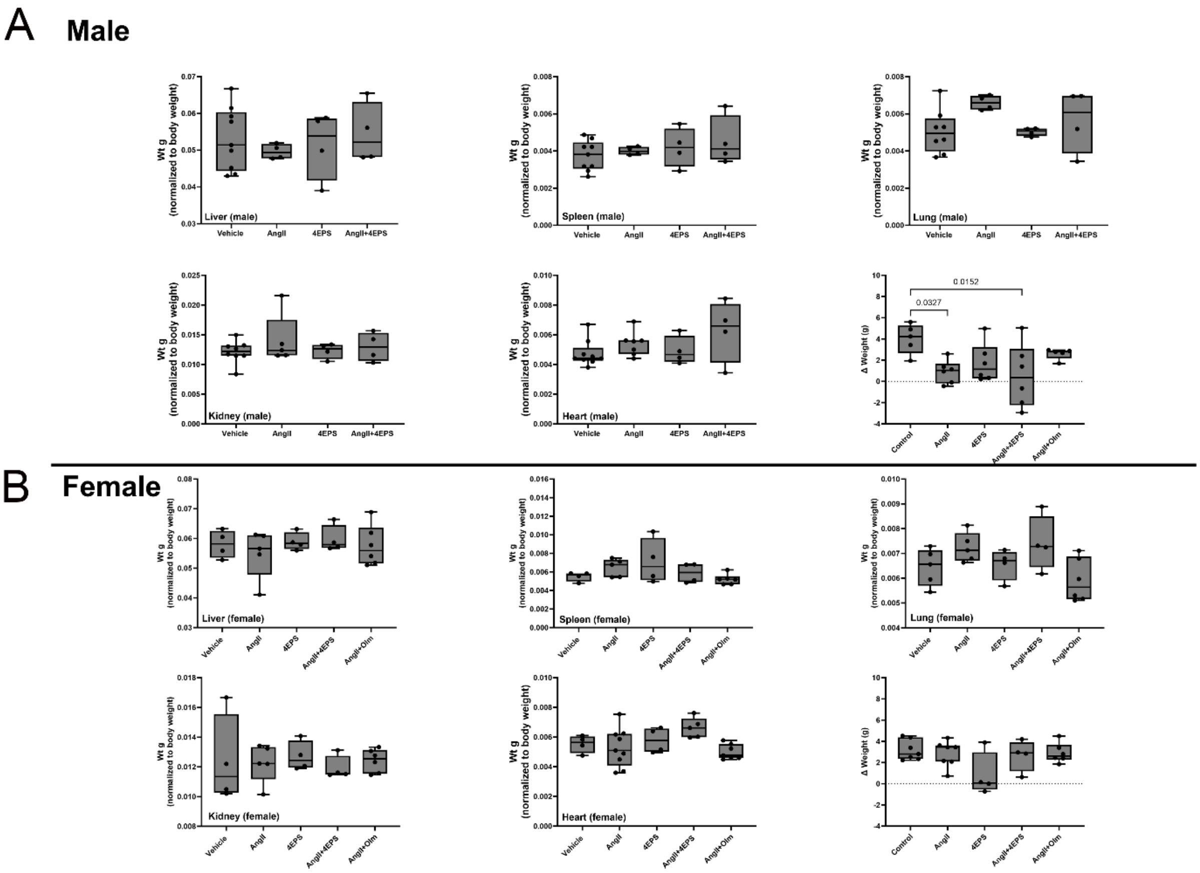


**Figure S3: Body weight and Organ weight measurements.** Organs harvested from both male and female ApoE-null mice on HFD infused for 28d with vehicle, AngII, 4EPS, AngII+4EPS and AngII+Olmesartan were assessed for changes is weight. Organ weights were normalized to total body weight and analysis was performed One way ANOVA with Dunnett's multiple comparison test using GraphPad Prism 10. N=4-12

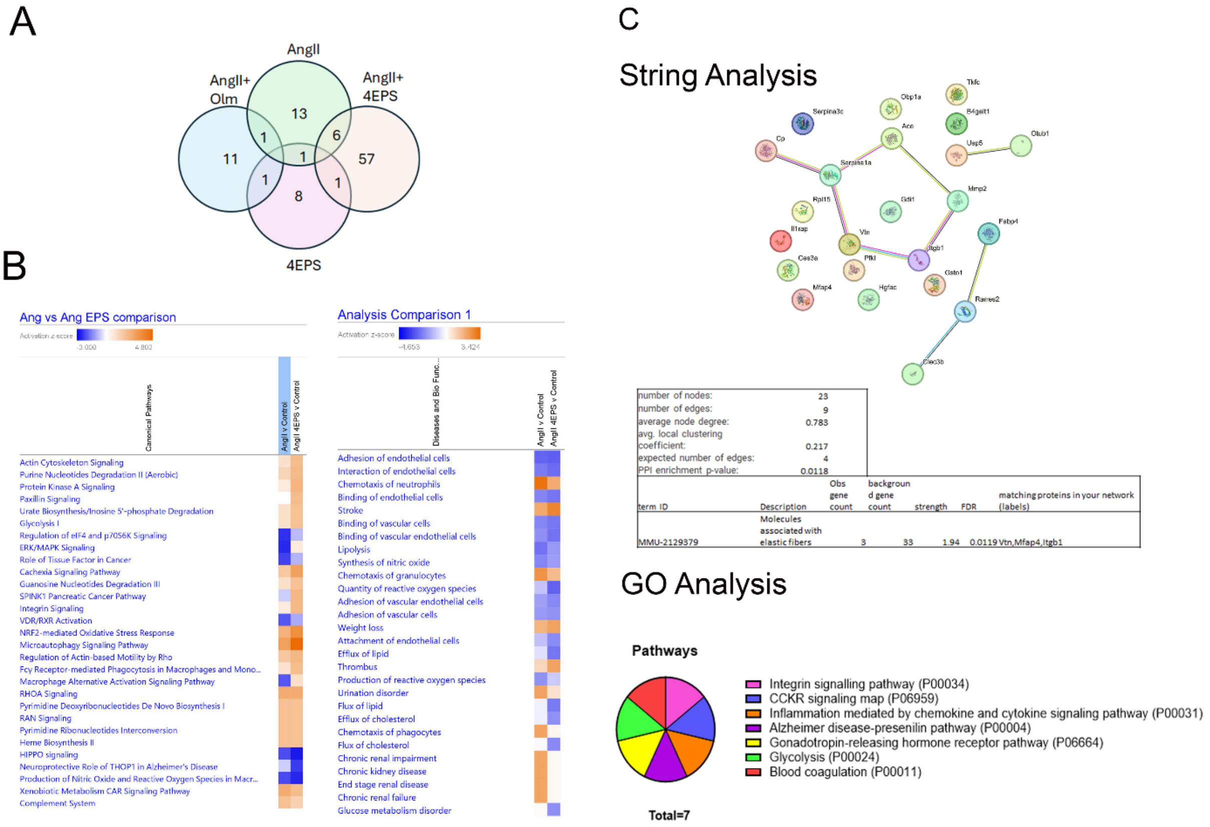


**Figure S4**: **Pathway analysis of proteomic data.** A. Venn diagram illustrating number of differentially expressed proteins between treatment groups. B Ingenuity Pathway analysis (IPA)(Qiagen) Top Canonical pathways (left) and Disease and Biofunctions (right) which were identified by IPA of proteins differentially expressed in plasma from AngII or AngII+4EPS infused ApoE-null mice. C. String database analysis of altered plasma proteins showing top reactome pathway being molecules associated with elastic fibers. D. Gene ontology database analysis identifying top pathways associated with altered plasma proteins.

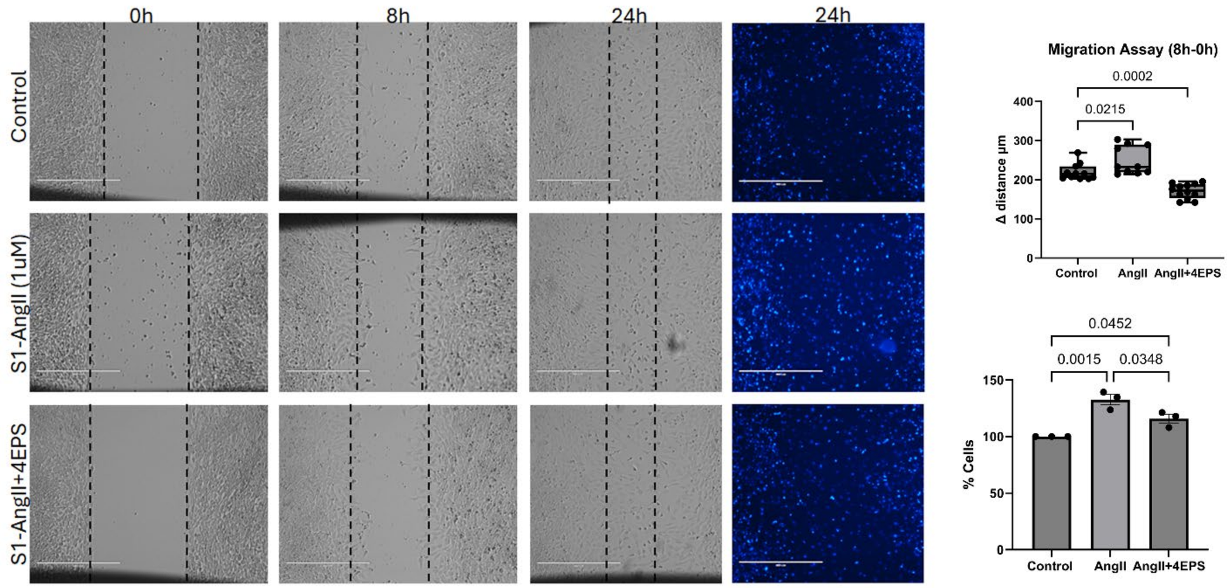


**Figure S5: Assessment of 4EPS on cellular migration of vascular smooth muscle cells.** MOVAS AT1R cells serum starved for 24hours, were treated with vehicle, [Sar1] AngII (10μM) or [Sar1]AngII +4EPS (100μM) and monitored for migration over 24 hours under serum free conditions. Images were acquired at 0, 8 and 24h, then stained with DAPI. After 24h, number of cells in the migrated zone were analyzed using ImageJ. Statistics were performed by One Way ANOVA with Tukey's multiple comparison test and normalized to Control =100%. Change in distance traveled Statistics were performed by One Way ANOVA with Dunnett's multiple comparison test. N=3 with 3 images per treatment per time point.

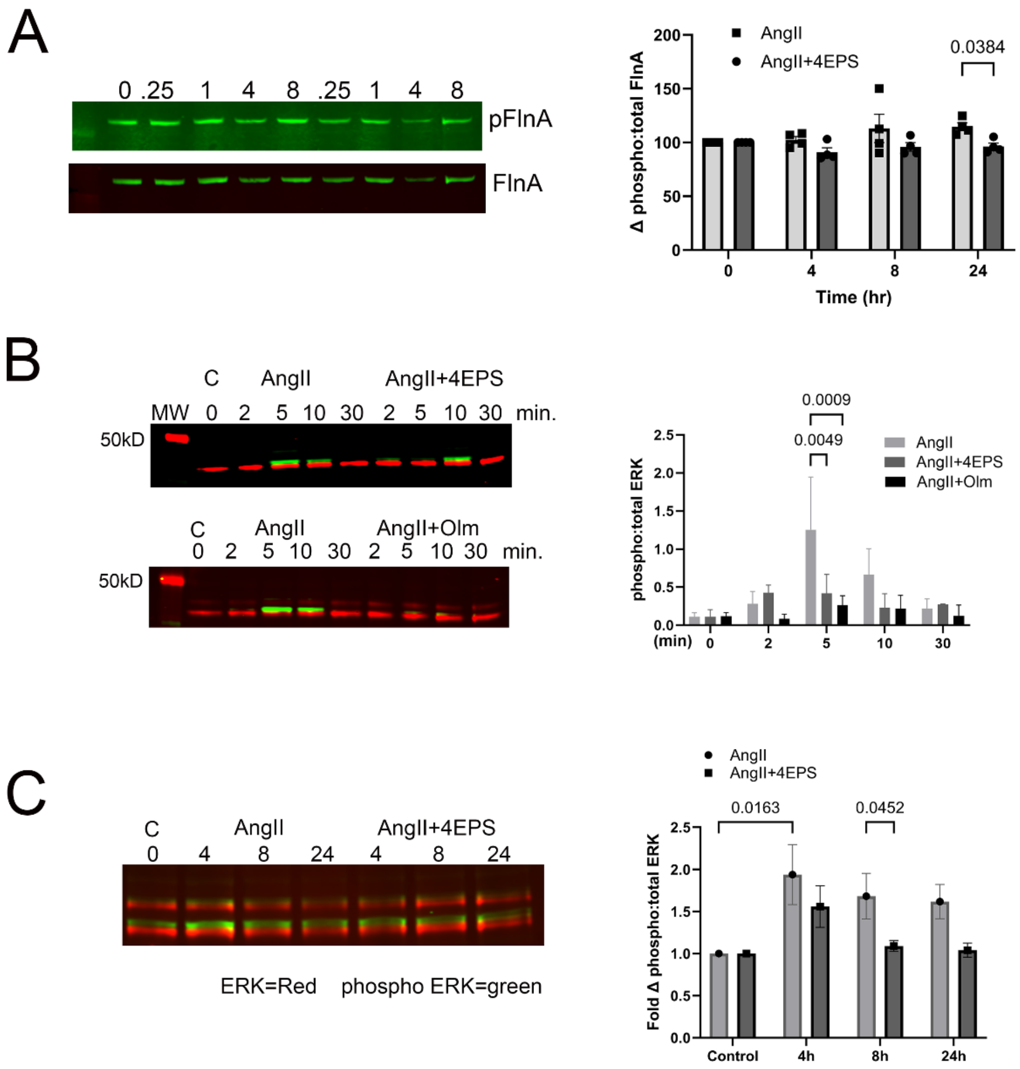


**Figure S6: Effect of 4EPS on Molecular Pathways**. Effect of AngII and 4EPS on expression and phosphorylation of FlnA and ERK in AT1R expressing cells. Immunoblotting analysis of expression and phosphorylation of A. FlnA in HEK-AT1R cells and B. MOVAS AT1R cells C. ERK (p42/44) in HEK AT1R cells and D. in MOVAS AT1R cells in response to AngII (4nM) or AngII+4EPS (100µM) at times indicated in figure. Phosphorylation of ERK or FlnA was normalized to total protein. Statistical comparisons were made at each time point between AngII vs AngII+4EPS N=3 and values expressed are mean ±SEM.

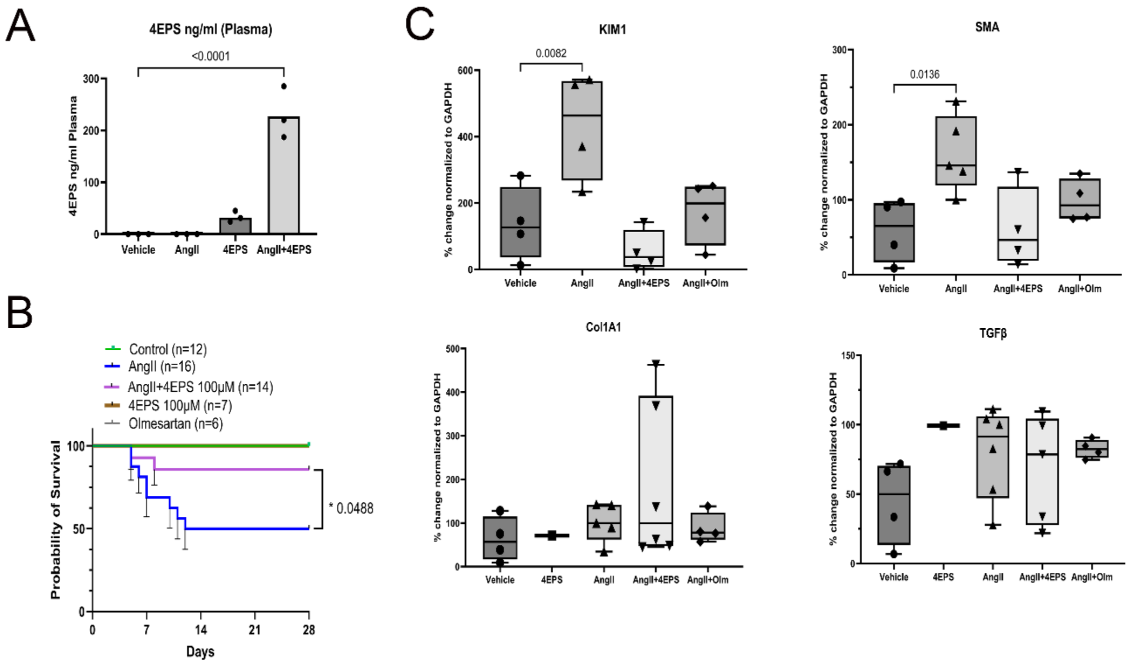


**Figure S7: Effect of 4EPS on survival and kidney function.** A. 4EPS levels were determined by mass spectroscopy in plasma from mice infused as described. B. Percent survival as represented by Kaplan-Meier curve were plotted at end of 28-day infusion or upon mortality as determined by aneurysm dissection seen during necropsy. C. Expression of KIM1 and SMA mRNA in kidney tissue homogenates from mice. Vehicle control was used as 100% as comparison for infusion groups.
